# Effective allogeneic natural killer cell therapy for pancreatic adenocarcinoma avails conserved activating receptors and evades HLA I-driven inhibition

**DOI:** 10.1101/2025.09.21.677560

**Authors:** Stacey N. Lee, Riley J. Arseneau, Thomas Arnason, Jeanette E. Boudreau

**Author notes:** Address for correspondence: Dr. Jeanette Boudreau, Department of Microbiology & Immunology, Dalhousie University, Halifax, Canada.

## Abstract

**Background:** At diagnosis, ∼80% of pancreatic ductal adenocarcinomas (PDAC) have metastasized. Relapse is thus common even among patients that undergo surgical resection, the only curative option. PDAC progresses rapidly, and existing immunotherapies have been ineffective. We hypothesized that natural killer (NK) cell immunotherapies could be effective against PDAC because they recognize conserved and heterogeneous features associated with cellular stress and transformation, and can seek out metastases distal to the primary tumour site. Here, we aim to define the key features of NK cells as effective agents for PDAC immunotherapy.

**Methods:** We used TCGA PDAC Firehose data and flow cytometry to predict and measure the most common activating or inhibitory ligands available on PDAC for NK cell activation. To ascertain how the tumour might alter expression of these ligands during treatment, inflammation or immune pressure, we measured expression of NK ligands at rest, or after exposure to immune cells or inflammation. To test and rank the functional importance of these dynamic ligands in the recognition, killing and control of PDAC< we used co-culture, antibody-blocking and an NK-competent humanized mouse model.

**Results:** Leveraging the known sequential acquisition of mutations as a surrogate for disease progression, we observed a progressive loss of transcript expression for activating NK cell ligands and chemoattractants. Exposure of PDAC to NK cells or IFN-γ, an inflammatory stimulus, drove dynamic changes in expression of both activating and inhibitory ligands. *In vitro* co-culture assays revealed a redundancy in the activating receptors engaged in NK:PDAC interactions, but that HLA-KIR signalling dominantly interrupted anti-PDAC activity. In NK-competent humanized mice, adoptively transferred, unselected, unmodified NK cells slowed tumour growth in a dose-dependent manner, but NK cells selected to avoid HLA I-driven inhibition were the most competent effectors for PDAC control.

**Conclusions:** Although there is redundancy among activating ligand:receptor pairs for recognizing PDAC tumours, but interactions between KIR and HLA define the extent to which anti-tumour activity can proceed. During tumour progression, and in response to immunotherapy. NK:tumour interactions drive upregulation of HLA I molecules. Thus, educated NK cells from HLA I-disparate donors may be the most effective allogeneic NK immunotherapy for PDAC.

**What is already known on this topic:** NK cells can be safely transferred across allogeneic barriers, so allogeneic therapy is possible. Since PDAC tumours progress very rapidly, there is insufficient time for engineered cell therapies. Although PDAC is typically considered to be an immunologically “cold” tumour, more recent studies have revealed that sub-tumour microenvironments can contain clusters of immune cells, and that the presence of immune cells in PDAC is associated with good prognosis.

**What this study adds:** We explore how the naturally-occurring heterogeneity of activating and inhibitory receptor expression with and between individuals impacts recognition of PDAC tumour cells. We find that the natural cytotoxicity receptors and NKG2D are among the most likely activating receptors to be expressed on NK cells responding to PDAC. However, interactions between inhibitory killer immunoglobulin-like receptors (KIR) and class I human leukocyte antigens (HLA) dominantly inhibit NK cell killing. To enable NK cell reactivity without the liability of inhibition, we demonstrate that NK cells can be selected intentionally from allogeneic HLA-mismatched donors, where they retain programmed functionality, but are ignorant to the HLA-driven signals for inhibition present on the tumour cells.

**How this study might affect research, practice or policy:** Diversity among NK cell functions is driven by interactions between HLA I molecules and KIR, with the remaining receptor-ligand partnerships mostly conserved across people. KIR:HLA I interactions are predictable, and there is a limited range of potential combinations, so definitions of key functions would empower mass-production of NK cells from a limited range of healthy donors that could be used as off-the-shelf cellular immunotherapy. Our study provides key exclusion criteria that will inform these selections.

**Lee Graphical abstract:** 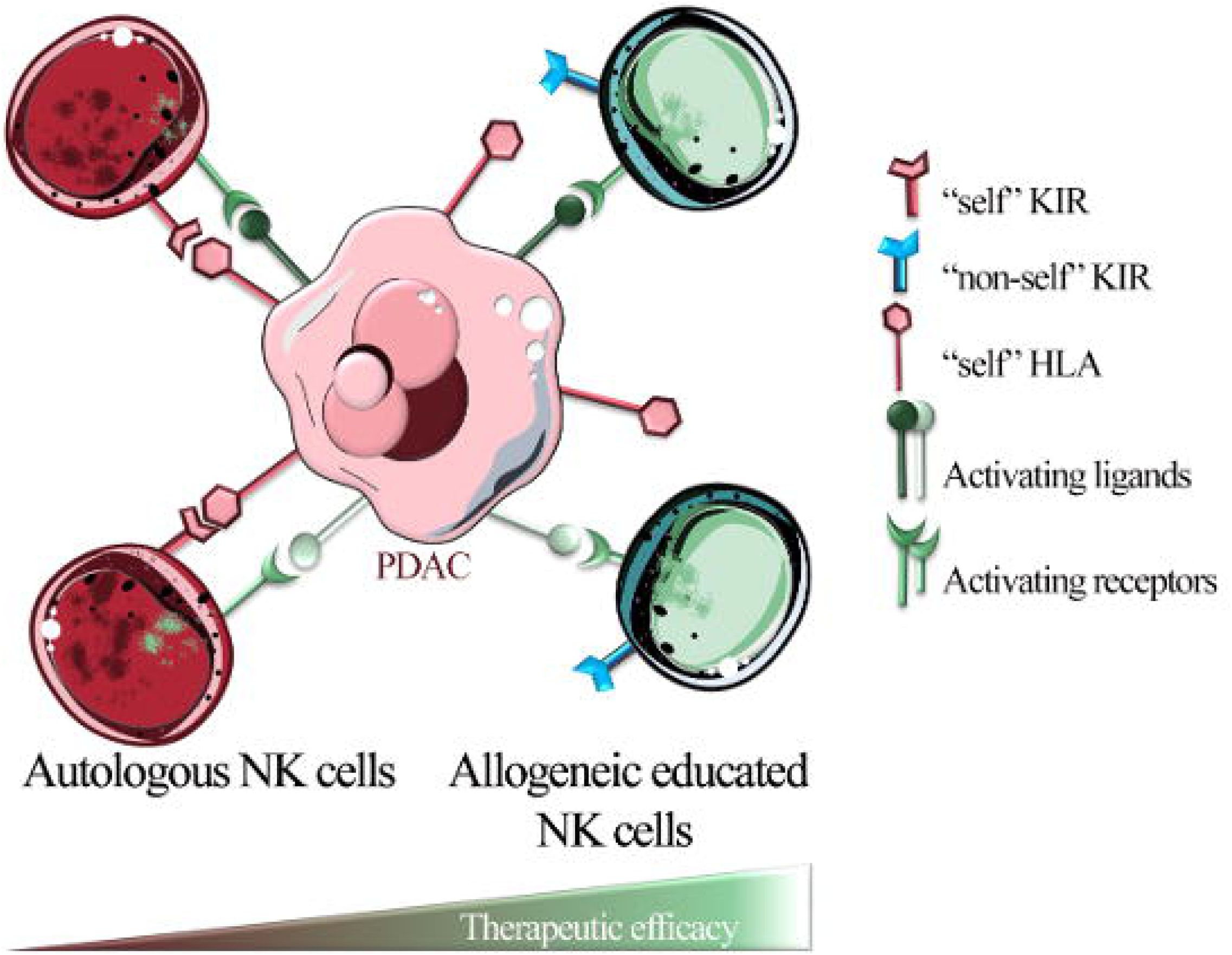

## INTRODUCTION

Pancreatic adenocarcinoma (PDAC) represents ∼90% of pancreatic tumours and is almost uniformly fatal without an aggressive pancreaticoduodenectomy (“Whipple”) procedure. Surgery is available to just ∼20% of patients: those diagnosed at stage II or earlier.^1^ Even among patients undergoing surgery, 60% relapse, revealing the presence of rapidly advancing, persistent and disseminated disease.^1^ One and five-year overall survival are 31% and 10%, respectively, making PDAC the 3^rd^ most frequent cause of cancer death, surpassed only by lung and colorectal cancers which occur 3-4 times more frequently.^2^ There is a clear and urgent need to define strategies to control PDAC at its primary and distal sites. Immunotherapy may be suitable for this purpose, but existing ones have not yet substantially increased survival among patients with PDAC.^3^ The ideal immunotherapy surveys the entire body, is available on demand, and recognizes the cancer’s phenotype as it mutates and spreads.

Clinical trials and pre-clinical studies demonstrate a susceptibility of PDAC to immune mechanisms, but also highlight that simultaneous targeting of multiple and dynamic features will be necessary for comprehensive tumour control. For example, a recent study revealed that mRNA vaccine-driven activation of T cells against cocktails of up to 20 patient-specific tumour-associated antigens with extensions of survival beyond three years.^4^ ^5^ PDAC is protected by its desmoplastic stroma, so other approaches have focused on improving immune cell assess to the tumour, such as with collagenase-armed CAR-T cells^6^ or oncolytic viruses^7^. Nevertheless, multiplex immunohistochemistry and genomic work have revealed that PDAC contains “sub” tumour microenvironments, many of which harbor immune cells, suggesting that immune-tumour interactions remain present^8^ ^9^ Indeed, infiltration and organization of immune cells into productive and multicellular neighbourhoods is associated with improved response to chemotherapy and overall survival.^10^ Thus, it is not the case that immune effectors are fully excluded from the tumour microenvironment and strategies that empower infiltration of well-armed effectors into PDAC might be most promising as future immunotherapies.

Natural killer (NK) cells recognize and kill targets that exhibit ligands upregulated consequent stress, transformation and DNA damage.^11^ Target cells – often derived from healthy “self” cells – can concurrently display ligands that signal to NK cells for activation or inhibition, but individual NK cells receive these signals differently because they have variegated expression and co-expression of germline-encoded receptors and thresholds for activation.^12^ ^13^ Inside the cell, NK cell receptors’ intracellular domains and adapters compete for intracellular kinases and phosphatases, so the net response of an NK-target interaction reflects the overall sum of incoming signals.^14^ ^15^ Each NK cell’s function is further calibrated by its ability to bind “self” class I human leukocyte antigens (HLA I), and thus differs between individuals.^14^ ^15^ This calibration (termed education, arming, licensing, or tuning) enables the hallmark “missing self” responsiveness of NK cells: killing of targets that lack “self” HLA I.^16^ ^17^ Though an educated NK cell is endowed with the strongest potential for activation by arming of additional activating receptors, HLA I binding to inhibitory receptors can override even strong signals.^18^ ^19^

The diverse and potent anticancer effects of NK cells make them ideal for cellular immunotherapy. Indeed, infiltration of solid tumours by NK cells is associated with beneficial prognoses^20^, even in patients with PDAC^21^. Strategies exist to modify NK cells with chimeric antigen receptors, expand NK cells virtually limitlessly, or induce adaptation with cytokines.^22–26^ NK cells can be safely transplanted across allogeneic barriers without graft-versus-host complications^27^, which makes it theoretically possible to have NK cells as on-demand therapy: a feature important in PDAC, a cancer that progresses very quickly.

We hypothesized that NK cells would be ideal effectors against PDAC if they could resist inhibition while continuing to respond to signals for activation. Using bioinformatics approaches, we found that primary PDAC tumours express NK cell ligands, especially those known to bind the activating NKG2D receptor, but expression of activating ligands decreases with the progressive accumulation of molecular changes. We find that the interaction between NK cells and PDAC drives alterations in the tumours’ expression of NK cell ligands. In particular, we observe a loss of activating ligands and concurrent upregulation of HLA I, which inhibited NK cell activation. In a humanized mouse model designed to support maintenance of NK cell function *in vivo*, we demonstrate that primary human NK cells control the growth of established PDAC tumours in a dose-dependent manner. When we selected HLA I-mismatched allogeneic NK cell donors to enable NK cell education dissociated from sensitivity to inhibition, tumour growth control was improved, underscoring an insensitivity to inhibition as key in the design of ideal NK cell-based immunotherapies for PDAC. Our findings imply that NK cell allogeneic therapy against PDAC will be improved by deliberate selection of HLA I-insensitive effector cells.

## MATERIALS AND METHODS

### PBMC and NK cell isolation

Peripheral blood mononuclear cells (PMBC) were isolated from buffy coats collected from healthy human donors via the Canadian Blood Services Blood4Research program. Use of PBMC was approved by the Dalhousie University REB (#2016-3842) and the Canadian Blood Services REB (#2016-016). In brief, human PBMCs were collected from buffy coats using Ficoll gradient centrifugation. Thereafter, primary NK cells were isolated using the NK Cell Isolation Kit (STEMCELL Technologies) per the manufacturer’s instructions. All PBMC donors were genotyped for *KIR* ligands (*HLA* subtypes with the HLA-C1, -C2 and/or Bw4 public ligands) and *KIR* genes using established primers and protocols.^28–30^

### Cell lines and IFN-**γ** treatment

The human pancreatic cancer cell lines AsPC-1 and PANC-1 (American Type Culture Collection (ATCC)) were cultured at 37°C in 5% CO_2_ in RPMI-1640 media with 10% heat inactivated FBS, 100 U/mL penicillin, and 100 mg/mL streptomycin. To assess effect of inflammation on cancer ligand expression and subsequent influence on NK cell activation, cell lines were treated with 1000 IU/mL of human recombinant IFN-γ (Peprotech) daily for three days prior to cocultures.

### TCGA analysis

RNASeq data was obtained from the Xena browser in October 2024^31^. RNASeq data is the gene-level transcription estimates, as in log2(x+1) transformed RSEM normalized count. RSEM normalization was performed by diving raw count values by the 75^th^ percentile (after removing zeros) and multiplying by 1000. Mutational data was downloaded from cBioportal “Type of Genetic Alterations Across All Samples” for genes of interest (*KRAS, TP53, SMAD4, CDKN2A*)^32^ ^33–35^. All non-benign small variants and copy number alterations (Amplifications, homozygous deletions) were included. Each point represents the expression of a gene for an NK cell ligand. The log2 fold change (mutated relative to wild type) was calculated, where positive values indicate higher expression in mutated samples and negative values indicate higher expression in wild-type samples. Empirical Bayes shrinkage was applied to improve variance estimates in small samples sizes. The p-value was Benjamini-Hochberg adjusted to account for the false discovery rate. Volcano plots were generated using Limma in R Studio ^36^.

### Co-culture and Flow cytometry

After thawing, PBMCs were rested overnight in RPMI with 100 IU/mL human IL-2 (Peprotech) at 1×10^6^ cells/mL. In a 96-well plate, PBMCs were co-cultured with AsPC-1 and PANC-1(3:1 ratio of effector:target cell) in the presence of anti-LAMP1 (CD107a, H4A3, BD Biosciences) for five hours at 37°C. All subsequent staining was performed at room temperature in the dark. Washing steps were performed by resuspending the cells in washing media to 200 mL (PBS or FACS wash buffer, depending on step) and spun at 300 x *g* for 5 min. Cells were stained with fixable viability dye V575 (BD Biosciences). To mitigate non-specific antibody binding, cells were incubated with 0.5mg/mL human Fc block (BD Biosciences) prior to antibody staining. Cells were then incubated with surface antibody staining cocktail diluted in Brilliant Staining Buffer Plus (BD Biosciences). Antibodies used are outlined in Supplementary Table 1-3. Finally, cells were fixed using 4% paraformaldehyde and resuspended in 200mL FACS wash buffer after washing.

FACS acquisition was performed using a BD FACS Symphony analyzer. To normalize acquisition metrics between experiments, we used 8-peak Rainbow Sphero beads (BD Symphony) and adjusted laser voltage to ensure equivalent mean fluorescence intensities across experiments. All exported files were compensated and analyzed using FlowJo v.10.7.1 (BD Biosciences). Viable NK cells were identified as CD56+CD3-with subsequent gating to identify key subpopulations (Supplementary Figure 1). Investigators were blinded to experimental conditions when appraising FACS data.

### Double PBMC:Tumour co-culture

AsPC-1 and PANC-1 cells were collected, counted, and plated in a 48-well at 1.85 x 10^5^ cells/well and then incubated at 37°C in 300 μL RPMI 12h prior to coculture to enable their adherence. For the first round of co-culture, PBMCs that had been rested overnight were pelleted, rinsed, and resuspended at 5×10^6^ cells/mL. From the pre-plated tumour cells, RPMI was removed and cells rinsed twice with PBS. PBMCs were then directly added to the wells with anti-CD107a (E:T 3:1) and incubated at 37°C for 10h. Thereafter, PBMCs were transferred to a 96-well plate for flow cytometry staining. Plated tumour cells were rinsed twice with PBS. PBMCs from the same (D1) or a different (D2) donor were then added to the plate for an additional 5h co-culture at 37°C. These “second round” PBMCs were transferred to 96-well plate for flow cytometry staining. Tumours cells were harvested by adding 100μL of trypsin to each well and incubated for 5 min. Tumours were then transferred to a 96-well plate, rinsed first with RPMI, then PBS,and stained for flow cytometry. Investigators were blinded to experimental conditions when appraising FACS data.

### Antibody blocking

Cancer cells were incubated human anti-HLA-ABC diluted in PBS (20 µg/mL, clone W6/32, ThermoFisher) for 30 min at 37°C. PBMCs were incubated with human Fc block for 30 min at 37°C. Cells were washed with PBS and resuspended for coculture and flow cytometry as described above.

### In vivo tumour models

Mouse experiments were approved by the Dalhousie University Committee on Animals (UCLA) (23-079 and 22-098), and adhered to the standards of the Canadian Council on Animal Care. Mice were group-housed in sterile conditions (2-5 mice/cage), with *ad libitum* access to autoclaved food and water and on a 12/12h light/dark cycle. All mouse experiments were performed using NOD/Rag^-/-^/IL2Rγ ^-/-^/Tg mice, which we backcrossed with NSG-Tg (Hu-IL15), NSG-HLA-A2 and NSG-HLA-Cw3 mice (all from the Jackson Labs) to create NRG mice that transgenically express human HLA I molecules. These represent the KIR ligand configuration present in AsPC-1 cells, which were used in all *in vivo* challenge experiments. Tumour cells were resuspended in 2X diluted Matrigel Matrix (Corning) and 1×10^6^ cells were injected into the left flank. Isolated human NK cells were intravenously injected 21 days post-tumour implantation (a time when tumours were palpable) at a dose of 1×10^6^, 2.5×10^6^, or 5×10^6^ cells. Mice were monitored and tumour volume measured using calipers every two days. Mice were randomly assigned to groups, and euthanized upon reaching humane endpoint and tumours were collected and fixed in formalin. Animals were selected to express the HLA I antigens HLA-A2 and HLA-Cw3 to match the AsPC-1 xenografted tumour cell line. No additional inclusion/exclusion criteria were employed.

### Hematoxylin and Eosin (H&E) staining

Tumours were fixed in 10% neutral buffered formalin, paraffin embedded (FFPE) and sectioned. Hematoxylin and eosin staining was performed and slides with imaged on a Mantra Quantitative Pathology workstation (Akoya) using the brightfield setting. Investigators were blinded to experimental conditions when appraising tissues.

### Statistical analysis

Expression data measured by flow cytometry show the mean ± standard deviation with significance indicated in the figure, unless otherwise specified. All statistical computations were performed using GraphPad Prism [Version 9]. Statistical significance was set at p<0.05, and significance levels are defined in figures as *, p<0.05; **, p<0.01, ***, p<0.001; ****, p<0.0001. Statistical tests and significance are also indicated within individual figure captions.

Normal distribution was assumed in experiments with eight donors or more. All co-culture experiments were completed in technical duplicate and the samples concatenated in FlowJo for downstream analysis. After concatenation, all duplicate samples were compared to ensure consistency between duplicates. In experiments comparing variation between >2 parameters with one independent variable, one-way analysis of variance (ANOVA) was completed with multiple comparisons. When comparing variation between >2 parameters with two independent variables ordinary two-way ANOVA with multiple comparisons was used. In the case where there was an uneven number of samples (such as when separating donors by sex), mixed-effects analysis was used with multiple comparisons. When comparing differences between two groups, nonparametric paired *t* tests were used. Sample sizes were chosen to enable power to detect statistically-significant differences at α=0.05.

## RESULTS

### NK cells infiltrate and kill PDAC *in vivo*

The function and education of NK cells differs substantially between mice and humans, so standard animal models are insufficient to inform human tumour-immune interactions. For this purpose, we created “autonomouse”, a NOD-RAG-IL2gc-/-mouse that transgenically expresses human HLA I and human IL-15 to support NK cell retention, proliferation, and education^37^ ^38^ (**Figure 1A**). To examine NK-tumour interactions *in vivo*, we bred autonomice to have the same KIR ligands as the PDAC cell line, AsPC-1 (i.e. HLA-C1). In them, we established subcutaneous PDAC tumours over three weeks, at which time tumours were approximately 150 mm^3^. Thereafter, we injected graded doses of NK cells from KIR ligand-matched healthy human donors (to mimic autologous NK cells) and monitored the progression of tumour growth (Figure 1A and B). At the highest dose of NK cells (5×10^6^), tumour growth stalled between weeks 4-7 before growth continuing (p<0.001). At the lower NK cell doses (2.5×10^6^ and 1×10^6^), tumour growth continued but was consistently slower than that of the control group, which received no NK cells (p<0.05). Although tumour growth was slowed by all three doses of NK cells, no dose eliminated or permanently stopped the growth of established PDAC.

**Figure 1.**
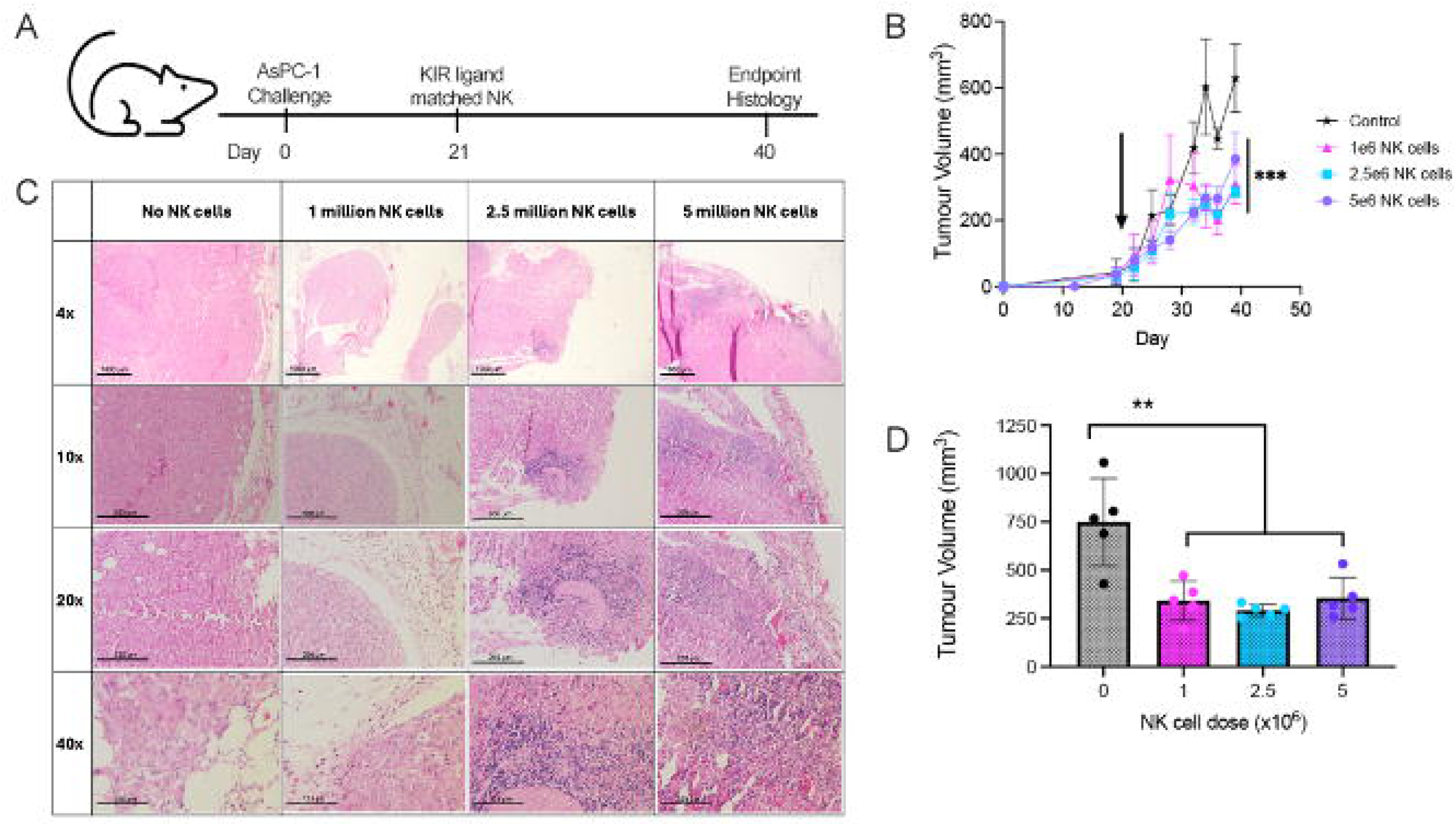
Human NK cells interfere with tumour growth *in vivo.* HLA and Hu-IL-15 transgenic mice were injected with 1 x 10^6^ AsPC-1 tumour cells and monitored for tumour growth. Three concentrations of unselected primary NK cells from healthy donor PBMC were injected intravenously 21 days after tumour implantation (arrow). Mice were monitored biweekly until the tumors reached a humane endpoint of 15mm along at least one length. (A) Methods for establishment of PDAC in our humanized mouse and experimental model; (B) Tumour growth with injection of PBS or increasing numbers of NK cells, measured with calipers, (***, p<0.001 compared with control); (C) H&E staining of FFPE tumors collected at endpoint, shown at different magnifications (Scale bars: 4x: 1000μm; 10x: 500μm; 20x: 250μm; 40x: 125μm). Note the accumulation of lymphocytes (purple) at the tumour’s margins, especially at the highest two doses. (D) Tumour volume at endpoint (**, p<0.01; all groups compared with endpoint). Data represents two experimental replicates with 3 mice per group per experiment (n=6).

Upon necropsy, tumours treated at any dose of NK cells were of comparable size (no NK: 749.7 ± 202.1 mm^3^; 1×10^6^ NK cells: 342.5 ± 90.6 mm^3^; 2.5 x 10^6^ NK cells: 291.5 ± 27.7 mm^3^; 5 x 10^6^ NK cells: 355.2 ± 95.1mm^3^). Tumour histology resembled that of malignant PDAC (**Figures 1C and D**). We confirmed lymphocyte infiltration into the tumour, especially at the periphery of tumours, with more present when mice had been treated with higher NK cell doses. H&E staining revealed alterations in structure of PDAC tumours with and without treatment with NK cells. Specifically, we note pyknosis – nuclear shrinkage – a hallmark sign of apoptosis, and a reduction in the density of tumour cells in tumours treated with NK cells. Immune cells are present along the periphery of tumour cells, especially at the highest two doses, implying an immune attack from the tumour’s edge and toward the middle of the tumours.

### PDAC displays NK cell activating ligands that decrease in response to progression or immune reactivity

Our humanized mice demonstrate that NK cells can interrupt growth of tumours *in vivo*. Despite this, even the highest dose of NK cells could not completely control tumour growth. We hypothesized that it may be possible to improve NK cell targeting of PDAC by selecting a subset of NK cells whose receptor combinations enable more precise recognition of PDAC cells. To understand which interactions might best enable NK cell targeting, we created a custom list of genes associated with NK cell activation or inhibition and used it to interrogate the TCGA Firehose database (**Supplementary Table 4**). PDAC tumours are understood to progress through a series of mutations^39^, starting most often with mutations causing *KRAS* overexpression (97.1% of PDAC), loss of *TP53* expression (74.8%), followed by mutations in *CDKN2A* (48.9%) and *SMAD4* (41.0%); thus, we compared expression of NK cell ligands between tumours with and without these mutations to emulate tumour progression (**Figure 2A-D**).

**Figure 2.**
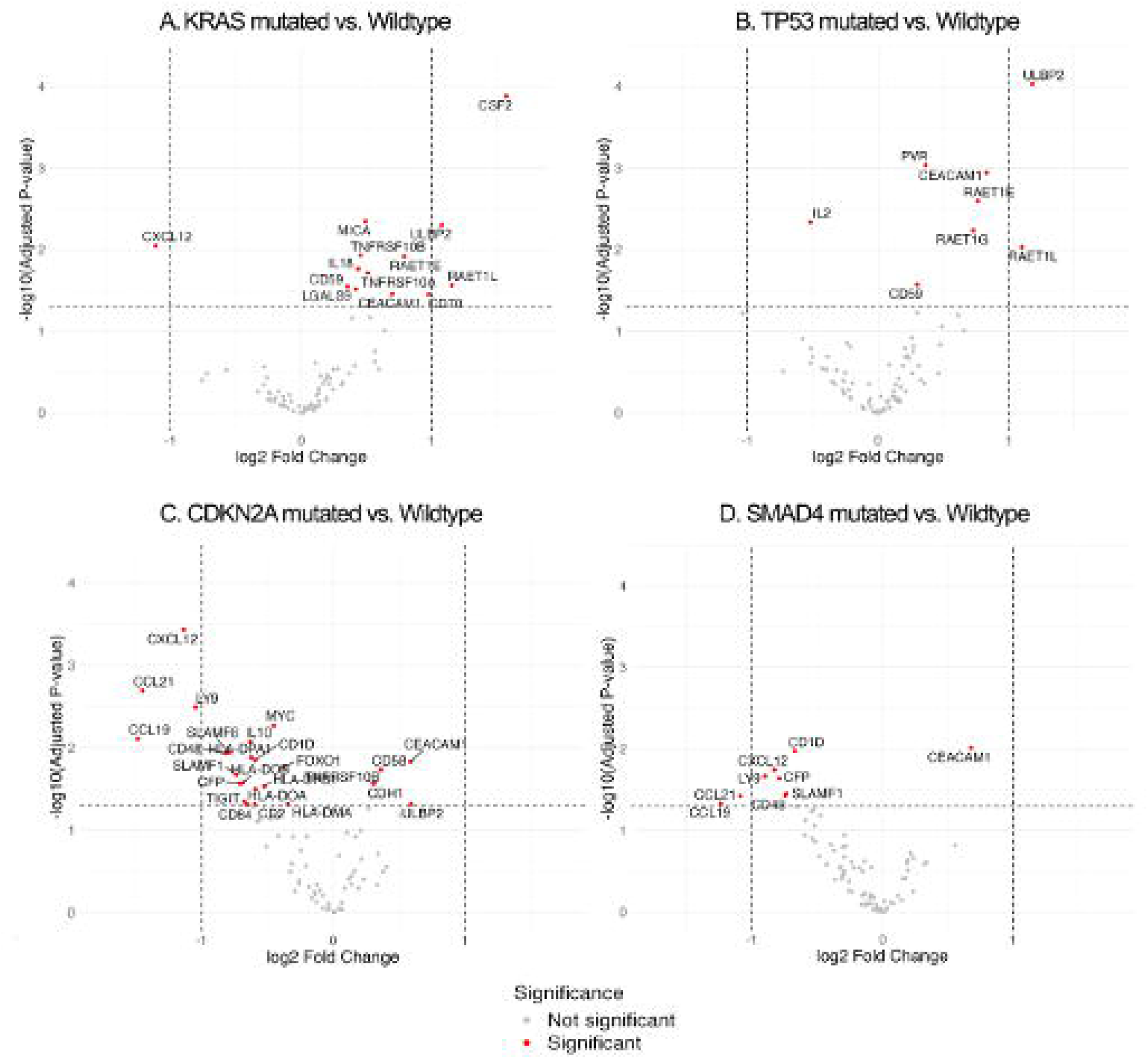
Differential gene expression of NK cell ligands in PDAC tumours from the Firehose Legacy cohort with genetic alterations in *KRAS, TP53, CDKN2A,* and *SMAD4*. Volcano plots showing differential gene expression of NK cell ligand genes in PDAC tumours (Firehose Legacy cohort^34^, accessed October 24, 2024) comparing patients with the indicated gene mutated to those with the wild type gene (n=147). Each point represents the expression of a gene for an NK cell ligand. The x-axis shows the log2 fold change (mutated relative to wild type), where positive values indicate higher expression in mutated samples and negative values indicate higher expression in wild-type samples. The y-axis shows the -log10 of the Benjamini-Hochberg adjusted p-values (False discovery rate). Horizontal dashed lines indicate the adjusted p-value threshold, and vertical dashed lines indicate the ±log2 fold chance cut-offs. Wild-type expression is used as the reference group in each comparison. **A)** *KRAS* mutated vs. wild type. **B)** *TP53* mutated vs. wild type. **C)** *CDKN2A* mutated vs. wild type. **D)** *SMAD4* mutated vs. wild type. Differential gene expression analysis was performed using the Limma R package, fitting a linear model to upper quartile-normalized RSEM RNA-seq data with empirical Bayers moderation.^56^

Nearly all PDAC tumours harbour gain-of-function mutations in *KRAS* oncogene and loss of function for the *TP53* tumour suppressor.^40^ Compared to tumours with wild type copies of these genes, those with non-benign mutations exhibited significant overexpression of genes for several ligands of the NKG2D receptor (i.e. *MICA*, *ULBP2*, *RAET1E*, and *RAET1E*; Figure 2A and B). Additional mutation in *CDKN2A* or *SMAD4* was associated with lower expression of NKG2D ligands, but a notable upregulation in the transcript for *CEACAM1*, which encodes a protein that binds with the immune checkpoint TIM-3, and has been associated with negative regulation of NKG2D.^41^ Similarly, in *KRAS*^mut^ tumours, there was an upregulation of galectin 9 (*LGALS9)*, another ligand for TIM-3, while *TP53*^mut^ tumours upregulated a *PVR*, which preferentially binds to the checkpoint receptor TIGIT.^42–44^ Thus, early mutations enabling PDAC (and disease progression) might allow for NK cell recognition, especially via the NKG2D (activating) receptor, but subsequent molecular alterations might enable escape from NK cell recognition via NKG2D.

### The expression of NK cell ligands on the tumour cell surface is dynamic, and changes in response to external stimuli

PDAC tumours are non-uniform in their genetic landscape^8^ ^10^ ^45^: progression is induced by chronic inflammation^46^ ^47^, and the engagement between NK cells and tumour cells might impact both populations. To consider the dynamic landscape of ligands expressed by tumours with which NK cells may interact, we created a flow cytometry panel that assesses ligand expression. We stained two PDAC cell lines for expression of NK cell ligands (**Figure 3**) (1) at rest; (2) after co-culture in the presence of PBMC from healthy donors; and (3) in response to IFN-γ. We selected two cell lines harbouring pathogenic molecular features associated with PDAC: AsPC-1 (*KRAS^G12D^TP53G*^404^*^T^CDKN2A^L78H^SMAD4^R^*^100^*^T^*) and PANC-1 (*KRAS^G12D^, TP53^R^*^273^*^H^CDKN2A^del^*).

**Figure 3.**
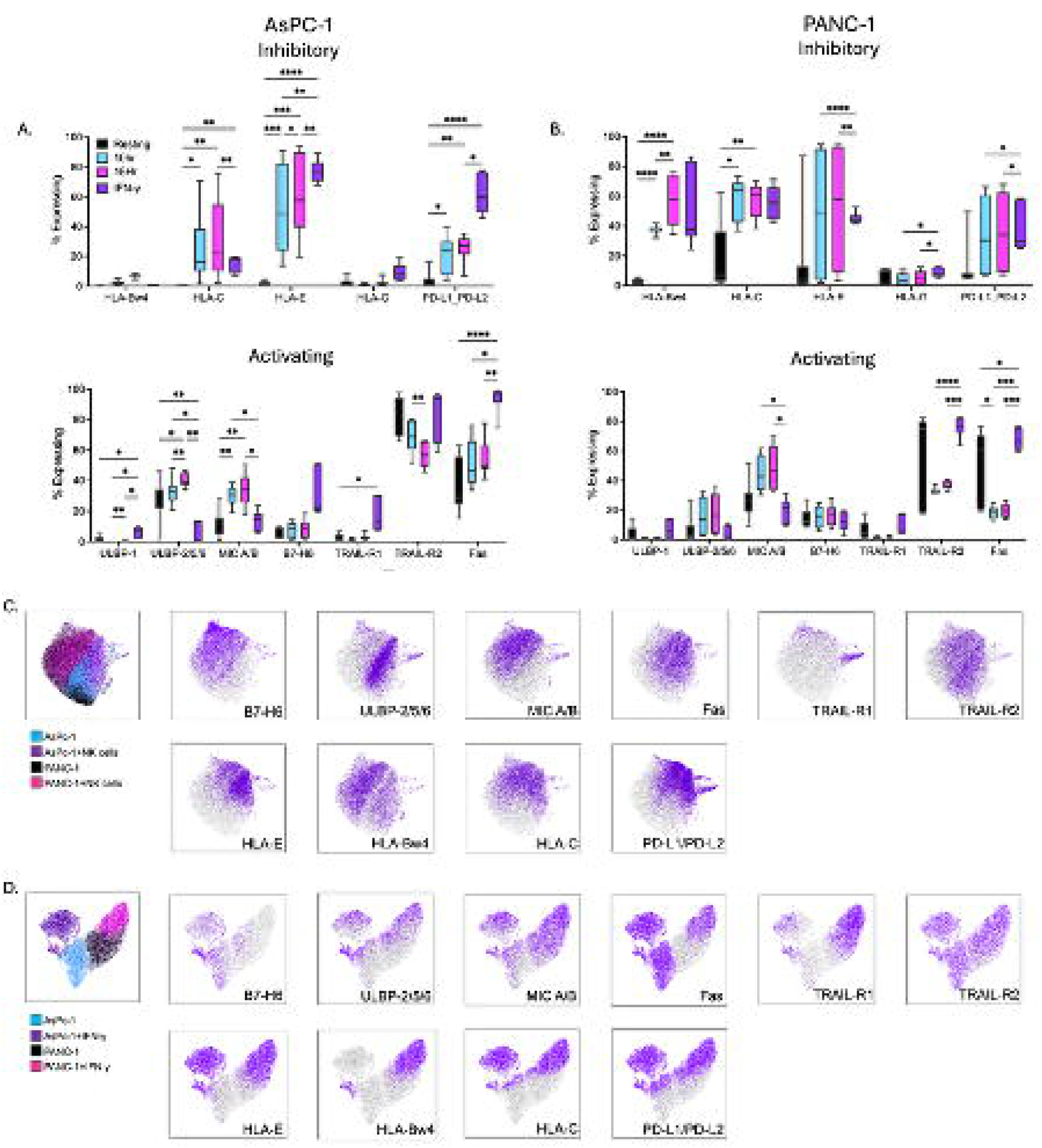
Inflammatory signaling results in increased expression of inhibitory cancer ligands and alters expression of activating ligands. (A, B) Ligands for NK cells were measured at rest, after a 10-hour co-culture with PBMCs (10h), a 15h coculture (with PBMC replaced at 10h) or after exposure to IFN-γ for 3 days. The percentage of tumour cells expressing each inhibitory (top) or activating (bottom) ligand is shown for each of (A) AsPC-1 (B) PANC-1 (n=12). (C, D) UMAPs depicting tumour cells’ ligand expression at rest or following exposure to (C) NK cells or IFN-γ (D) Data are representative of 16 independent donors or 6 trials. *=p < 0.05; **=p<0.01; ***= p < 0.001; ****=p<0.0001

Almost universally on both cell lines, PBMC co-culture and IFN-γ exposure induced expression of inhibitory ligands (**Figure 3A and B**). At rest, HLA and PD-L1/L2 were very poorly expressed. For AsPC-1, expression of HLA-C was highest after co-culture with primary PBMCs (30.6%±25.3%), though this expression was more variable and donor dependent, compared to expression induced by IFN-γ (15.2%±5.6%). This was also true for expression of HLA-E by AsPC-1. HLA-C expression on PANC-1 was similar across the treatment groups (PBMC: 57.5%±10.6%; IFN-γ: 55.7%±12.1%). AsPC-1 do not express a gene any HLA I encoding the Bw4 ligand; unsurprisingly, it was not induced on this cell line. HLA-Bw4 was, however, upregulated on PANC-1, which does encode this gene. Finally, the expression of PD-L1/L2 was most expressed by the IFN-γ treated AsPC-1 cells (61.5%±13.5%) in comparison to those exposed to donor PBMCs (25.1%±8.4%). Altogether, these findings indicate a persistent upregulation of inhibitory NK cell ligands, as tumours respond to inflammation; these changes would be expected to interrupt NK anti-tumour functions.

At rest, the phenotypes of AsPC-1 and PANC-1 were similar, but co-culture with PBMCs or the addition of IFN-γ to incite inflammatory responses diversified the phenotypes of the two cell lines (**Figure 3A and B**). Ligands for the activating receptors FAS-L (Fas; AsPC-1:36.8%±17.3%; PANC-1:48.5%±25%) and TRAIL (TRAIL-R2; AsPC-1: 84.3%±13.8%; PANC-1: 56.6%±29.4%) were present on both cell lines at rest and their expression was retained or enhanced in the presence of IFN-γ (Fas; AsPC-1:93.9%±9.45%; PANC-1: 67.5%±7.3%. TRAIL-R2; AsPC-1:84.2%±16.9%; PANC-1:75.9%±6.9%). Exposure of these same tumour cells to primary PBMCs led to the decrease of expression of TRAIL-R2 on AsPC-1 (56.8%±9%), though expression was retained on PANC-1.

Some expression of ligands for the activating receptor NKG2D was observed at rest, but their expression was dynamic in response to inflammatory and immune pressures. Specifically, ULBP-2/5/6 was favoured in AsPC-1 (25.5%±13.8%) and MICA/B was favoured in PANC-1 (26.1%±12.1%). Both activating ligands were reduced by exposure to IFN-γ (ULBP-2/5/6: 8.9%±6.6; MICA/B:19.5%±8.4%). In response to PBMC coculture, expression of both ULBP-2/5/6 (40.1%±4.2%) and MICA/B (33.3%±9.6%) was significantly elevated on AsPC-1 and trended similarly in PANC-1. B7-H6, a activating ligand for the natural cytotoxicity receptor NKp30, was consistently present on a PANC-1 (14.6% ± 6.5%) cells with and without IFN-γ stimulation (12.8%±6.6%) or co-culture with PBMCs (16.2%±7%). AsPC-1, expressed low B7-H6 at rest (4.8%±3.6%), but this ligand was upregulated in response to IFN-γ (32.4%±16.5%) or after PBMC co-culture (8.2%±5.2%). CD112/PVR, which binds both DNAM-1 and TIGIT, was only available after exposure to IFN-γ, but was almost universally expressed on both cell lines with and without treatment (**Supplementary Figure 2**). Taken together, bioinformatics assessments and tumour cell staining *in vitro* are both consistent with the dynamic and heterogeneous availability of activating and inhibitory ligands for NK cells, supporting strategies that leverage NK cells with an array of cognate activating receptors but lacking inhibitory receptors that could concurrently bind tumours and ultimately interrupt signals for activation.

To ascertain the combinations of inhibitory and activating ligands present on individual tumour cells (and thus infer the ideal configuration of NK cell receptors), we used Uniform Manifold Approximation and Projection (UMAP) analysis to understand co-expression. Cells were examined at rest or following exposure to NK cells or IFN-γ (**Figure 3C and D, Supplementary Tables 2 and 3**). PDAC cell lines were similar to each other at rest, but patterns of ligand expression distinguished them following inflammatory pressures. We observe a redundancy of both activating and inhibitory ligands on both cell lines after IFN-γ or NK cell exposure, revealing that NK cells are likely to concurrently receive both signals from target PDAC cells.

### NK cells’ responsiveness is reduced by inhibitory ligands

Bioinformatics, tumour ligand phenotyping, and co-culture analysis each revealed dynamism among PDAC ligand expression, which we predicted would impact the reactivity of NK cells against PDAC. Against a tumour in a patient, the stages of inflammation, rest, and immune pressure would be all expected to be present and dynamic, and effective NK cell therapy would need to respond to these conditions. To discern which signals might dominate in the inflammatory tumour environment, we co-cultured NK cells with our PDAC cell lines before and after IFN-γ stimulation (**Supplementary Figure 3**). NK cell responsiveness was reduced after this inflammation, despite upregulation of activating ligands, suggesting that co-expressed inhibitory ligands dominate over concurrent signals for activation.

We next measured the phenotypes of NK cells responding to tumour cells under the different conditions of inflammation and NK cell pressure. As above, we cocultured PBMCs and tumour cells for 10hr, washed, and then re-cultured with PBMCs from the same (D1) or a different (D2) donor for an additional 5hr. In this way, we could understand whether induced changes in tumour phenotypes were universally inhibiting to NK cell function and using flow cytometry-based phenotyping, we could directly measure the receptors involved in NK cell responses (**Figure 4A-C**).

**Figure 4.**
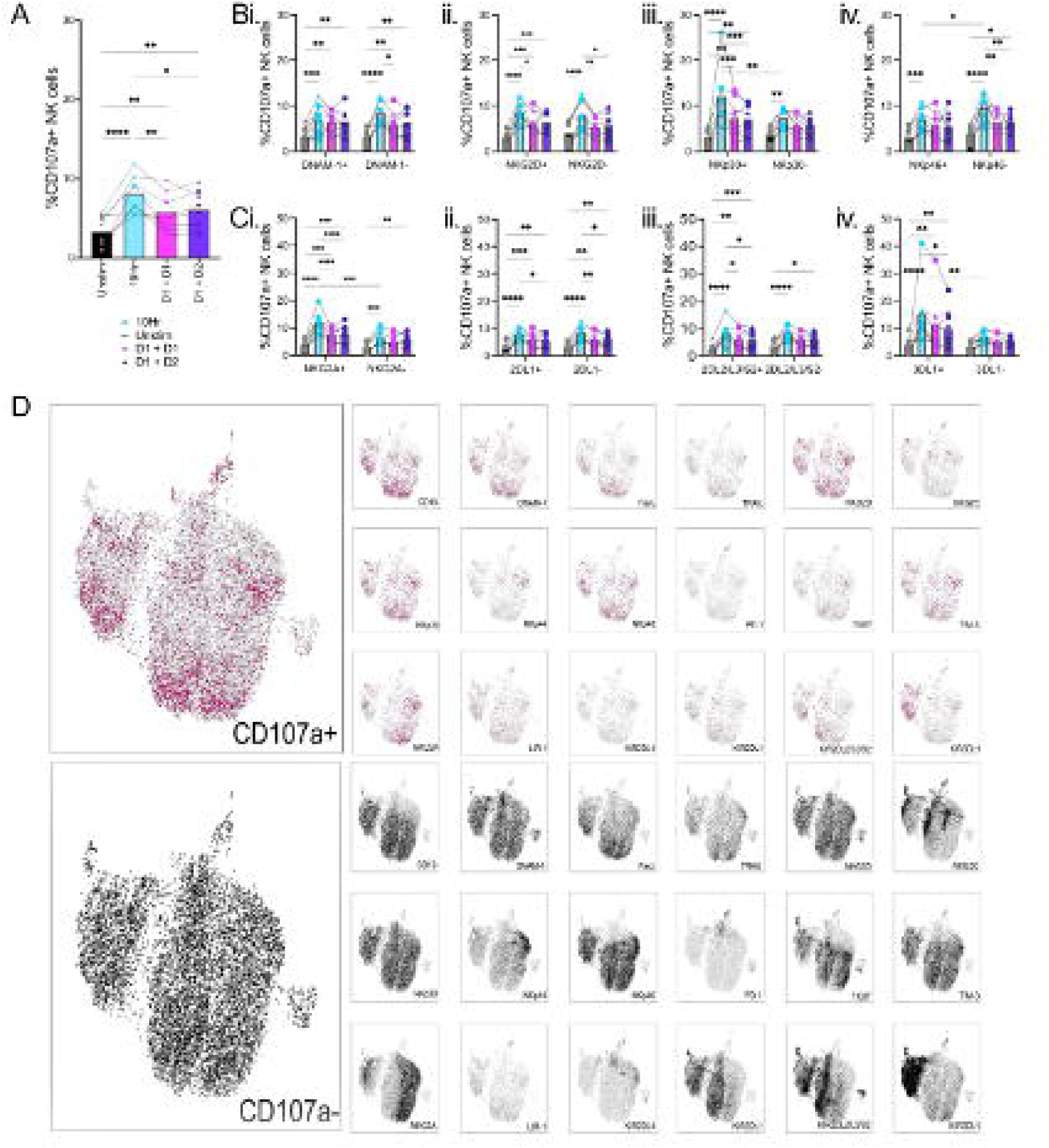
Activation induced tumour ligand changes reduce responsiveness of primary NK cells. To assess how the tumour ligands’ changes impact the subsequent responsiveness of primary NK cells, PBMCs were co-culture with the PDAC cell line AsPC-1 for 10-hour (10 Hr), then rinsed and re-co-cultured for 5-hour, with PBMCs of either the same donor (D1+D1) or a donor with a different educational background (D1 +D2) (A) Changes in total NK cell degranulation after 10hr or 10hr + 5hr co-culture with AsPC-1 (B) Degranulation of activating receptor positive of negative populations (i. DNAM-1, ii. NKG2D, iii. NKp30, iv. NKp46). (C) Degranulation of KIR-or KIR+ populations (i. NKG2A, ii. KIR2DL1, iii. KIR2DL2_L3_S2, iv. KIR3DL1). (D) UMAP of responding (pink) versus non-responding (black) NK cells after co-culture with AsPC-1. Overlays of each receptor population is shown (n=8 donors). Statistics shown are two-way ANOVA with multiple comparisons; *=p < 0.05; **=p<0.01; ***= p < 0.001; ****=p<0.0001

Degranulation of NK cells against AsPC-1 was highest among NK cell during the first 10hr of culture for all donors, consistent with low expression of inhibitory ligands on resting tumour cells (**Figure 4A**). Regardless of whether NK cells were from the same or a different donor, NK cell responses were universally lower during the second stimulation, after the targets had already changed or been selected by NK cells in the first 10h. To ascertain which receptor(s) drive responsiveness or inhibition in response to PDAC cells, we drilled down on cognate activating receptor+ populations for the ligands known to be expressed or induced by PDAC cells (**Figures 2 and 3**). These included the activating NKG2D, NKp30, NKp46 or DNAM-1. Only NKp30 expression (but not NKG2D or DNAM-1) was associated with increased responsiveness to AsPC-1 target cells during the 10hr co-culture (**Figure 4B**). Against PANC-1, it was instead DNAM-1 expression that best marked responding cells (**Supplementary Figure 4**). These responses were diminished across all donors in the second round, suggesting that the pre-conditioning or prior exposure of tumour cells to NK cells induces changes consistent with escape from NK cell-mediated recognition.

Expression of activating and inhibitory receptors is co-incident, and NK cells that express KIR and/or are capable of “missing self” responsiveness (i.e. educated) arm more activating receptors than their KIR-negative/uneducated counterparts and express greater densities of cytotoxic granzymes.^48^ ^49^ We therefore next included expression of NKG2A or KIR to determine whether these receptors might tune NK cell responsiveness. Against AsPC-1, NK cells expressing NKG2A and KIR3DL1 exhibited greater responsiveness than cells lacking these receptors during the first 10hr of co-culture (**Figure 4C**). However, no significant differences were observed as a function of NKG2A or KIR against PANC-1 (**Supplementary Figure 4**).

The enhanced responsiveness available to KIR/NKG2A+ NK cells is a result of their education to respond to “self” HLA I molecules. Hence, the upregulation of HLA I might interrupt activating signals. To explore this hypothesis, we again used UMAPs to visualize how NK cell phenotypes might be enriched or underrepresented among responding populations (i.e. CD107a+ vs. CD107a-) (**Figure 4D**). Each of DNAM-1, NKG2D, NKp30 and NKp46 were present among responding subpopulations of NK cells, consistent with their widespread and conserved use throughout the NK cell repertoire. While these included populations that co-express inhibitory NKG2A and KIR, we note that these receptors are generally underrepresented among responding populations. Noteworthy, however, NK cells expressing the inhibitory KIR3DL1, whose ligand is not expressed by AsPC-1 cells, remained among the responding subtypes. From this, we conclude that inhibition of NK cells via cognate HLA-KIR/NKG2A interactions may be the principal factor interfering with activation in NK:PDAC interactions.

### Induced inhibition dominantly interrupts NK cell responsiveness

To formally test the role of HLA-KIR binding as a driver in interrupting NK cell degranulation against PDAC, both resting and IFN-γ pre-treated AsPC-1 and PANC-1 were incubated with a pan anti-HLA antibody that blocks HLA-A, B, C, and E from interacting with KIR.^50^ ^51^ In this context, NK cell NK cell activation was restored, confirming this pathway as the dominant one interfering with NK cell activation (Figure 5A and **Supplementary Figure 5**). Against AsPC-1, which does not express HLA I at rest (Figure 3A), HLA blocking restored NK cell activation targeting the IFN-γ pretreated cells. PANC-1 does express low levels of HLA I at rest, though it is also upregulated after exposure to IFN-γ (**Figure 3B**); blocking both restored responsiveness against IFN-γ treated cells, and enhanced responsiveness against resting PANC-1 (**Supplementary** Figure 5A).

**Figure 5.**
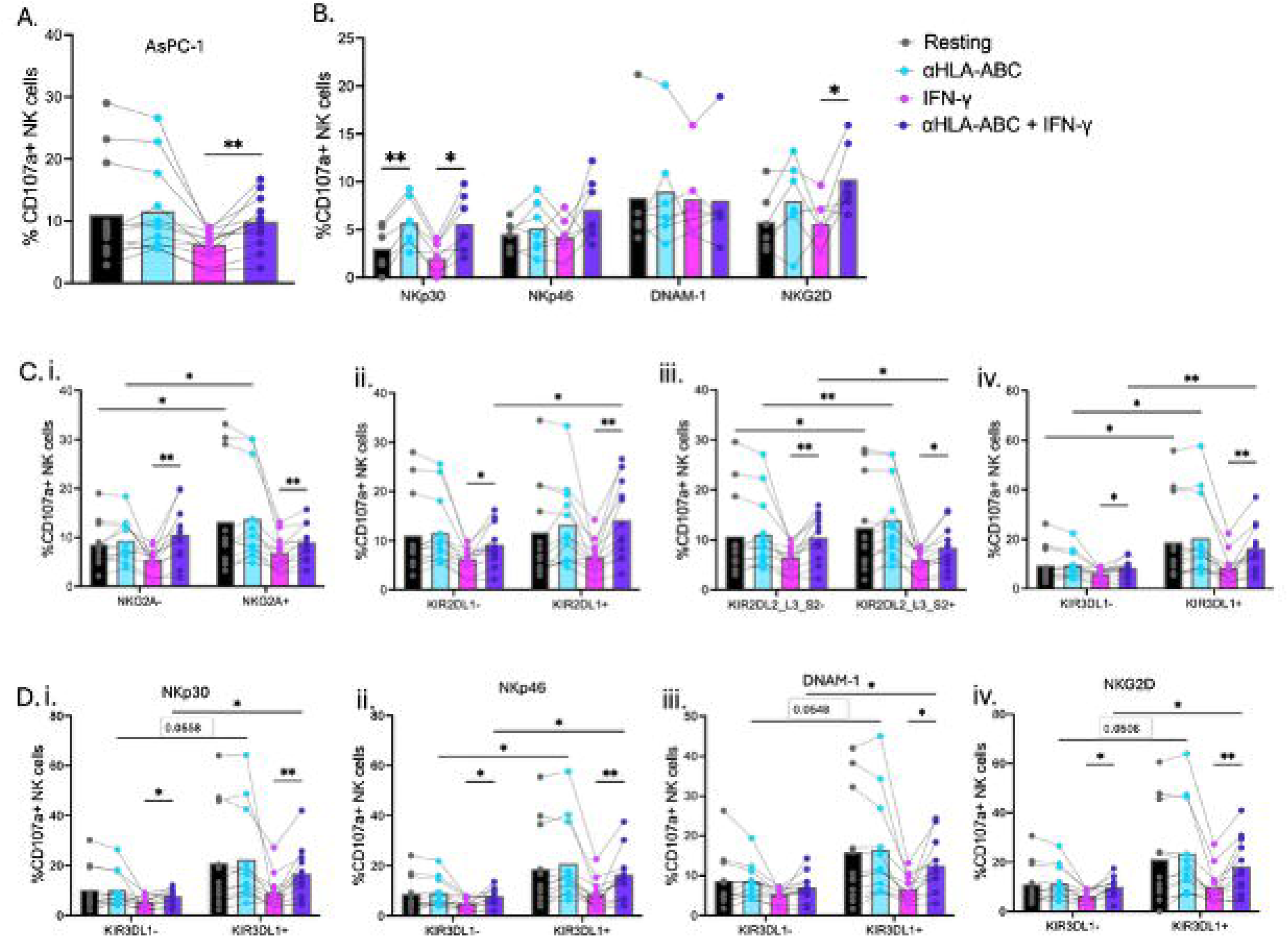
HLA I inhibits NK cell reactivity against PDAC tumour cells. AsPC-1 tumour cells were treated with IFN-γ or left untreated for three days, prior to co-culture with primary human NK cells (3:1 E:T) in the presence or absence of HLA I blocking antibody (αHLA) to identify the impact of HLA-driven inhibition on NK reactivity against PDAC. (A) changes in total NK cell degranulation in response to AsPC-1 treated with IFN-γ and/or HLA-ABC blocking after 5h of coculture. (B) Degranulation of single positive activating receptors (C) Degranulation of KIR- or KIR+ populations (i. NKG2A, ii. KIR2DL1, iii. KIR2DL2_L3_S2, or iv. KIR3DL1). (D) Activating receptor degranulation when co-expressing, or not, KIR3DL1 (i. NKp30, ii. NKp46, iii. DNAM-1, iv. NKG2D). Lines join the responses of individual donors (n=12). *=p<0.05; **=p<0.01

Seeking to define the key activating receptors involved in PDAC recognition, we drilled down on NK cells co-expressing receptors until only the single positive populations remained. Against AsPC-1, only NKp30+ NK cells were significantly rescued in the presence of HLA blocking (**Figure 5B**). For NK cells responding to PANC-1, NKG2D was best associated with responsiveness when HLA I-mediated signaling was blocked, with NKp30^+^ NK cells trending similarly (**Supplementary Figure 5B**). Although these reveal NKG2D and NKp30 as contributing receptors in PDAC recognition, we note that other activating receptor expressing NK cells are nevertheless recruited for activation against PDAC, suggesting a putative redundancy among them.

To discern whether KIR and NKG2A (and their associated inhibition) dominate NK cell responsiveness against PDAC, we compared their responsiveness with and without blocking (Figure 5C). KIR and NKG2A positive populations were indeed more responsive than the negative populations, with both KIR2LD2/L3 and KIR3DL1 expressing cells associated with increased activation against AsPC-1 in both resting and IFN-γ HLA I-blocked conditions. Against PANC-1, it was the KIR3DL1^+^ NK cells that were higher responders in both conditions. However, when we assessed how the co-expression of these KIR populations impacted the responsiveness of each activating receptor, only KIR3DL1 expression was correlated with increased activation when HLA I interactions were blocked in both conditions (**Figure 5 D; Supplementary Figure 5 and 6**), and rescued activation among NKp30, NKp46 and NKG2D-expressing NK cells when they were co-expressed with KIR3DL1 (**Figure 5D**) with similar results observed against the PANC-1 cell line (**Supplementary Figure 5D**). Taken together, these experiments reveal KIR-HLA interactions as important checkpoints interfering with NK cell recognition of PDAC and as the key considerations in the design of NK cell-based immunotherapies.

### KIR+ NK cells permit continued and superior control of PDAC in vivo

Our results demonstrate that HLA-driven inhibition is the dominant factor defining NK cell responsiveness against PDAC, but activating receptors have some redundancy for recognition. Since a patient’s NK cells are educated by the same autologous HLA I molecules expressed by their tumours, HLA I-driven inhibition will interfere with productive NK-driven cancer control. Allogeneic NK cells, however, could offer the opportunity to dissociate sensitivity to inhibition from NK cell activating receptors if donors are selected intentionally to avoid this inhibition. To test this hypothesis, we repeated our “autonomouse” xenograft studies, but selected NK cell donors whose KIR ligands matched (HLA-C1) or did not match (HLA-Bw4) the ligands that could be expressed by AsPC-1, our model PDAC *in vivo* (**Figure 6A**). Indeed, NK cells from donors lacking the same KIR ligands as the tumour were superior effectors of tumour control compared with those whose ligands matched those of the tumour (**Figure 6B and D**). Upon necropsy, tumours treated at with HLA-Bw4 educated NK cells were small than the uneducated, HLA-C1 only NK cells (HLA-Bw4: 141.8±49.3 mm^3^; HLA-C1:236.1±47 mm^3^), though either donor had decreased tumour growth compared to the no NK cell control (417.2±82.7 mm^3^). There was immune infiltration into the tumours treated by either donor, especially at the periphery of tumours as noted in the earlier study (**Figure 1C and 6C**). Taken together, our findings support the use of allogeneic NK cells for adoptive cell therapy of PDAC and reveal HLA-driven inhibition as a dominant, but avoidable, pitfall to the success of this therapy.

**Figure 6.**
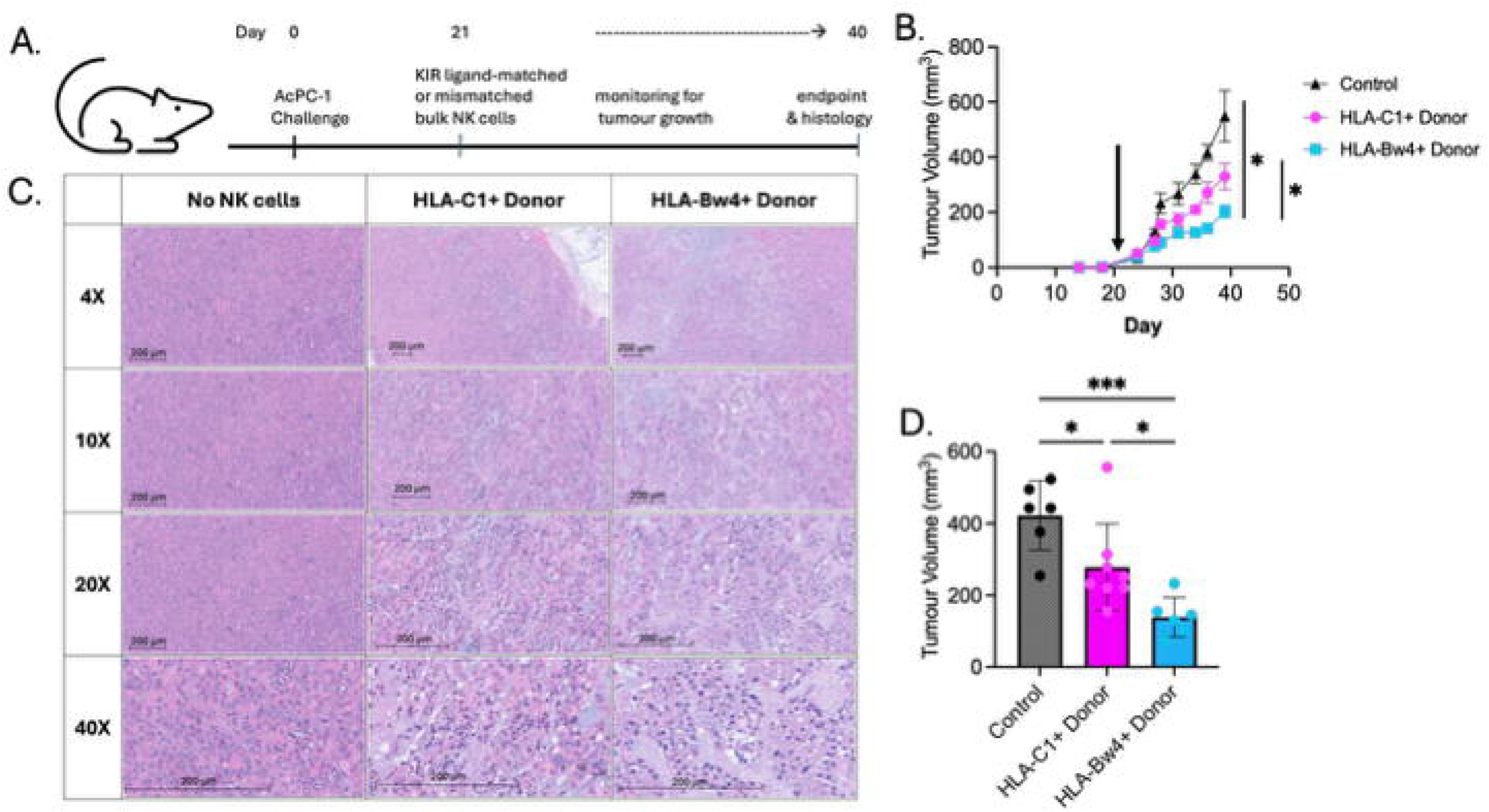
NK cells educated by non-self HLA ligands are more effective at tumour growth control *in vivo*. Our humanized mice (NOD/RAG1^-/-^IL2Rγc^-/-^ IL-15^+/+^ HLA-C1 transgenic) were injected with 1 x 10^6^ AsPC-1 tumour cells (HLA-C1+/ HLA-Bw4-) and monitored for tumour growth. NK cells from healthy donor PBMCs (5×10^6^ cells) derived from an HLA-Bw4 educated donor (non-self) or HLA-C1 (self) donor were injected intravenously 21 days after tumour implantation (arrow). (A) experimental protocol (B) Tumour growth with injection of PBS or NK cells of different educational background, measured with calipers, (*=p<0.05 compared with HLA-C1 treated tumour and control). (C) H&E staining of FFPE tumors collected at endpoint, shown at different magnifications (scale bar set at 200μM). (D) Tumour volume at endpoint. *=p<0.05; ***=p<0.001. Data represents two experimental replicates with 4 mice per group per experiment (n=8/group total).

## DISCUSSION

NK cells are garnering justifiable optimism for use as cellular therapies. Protocols exist for the nearly limitless expansion of cells *ex vivo*, cytokine treatment to induce longevity and potent effector function, or engineering for expression of chimeric antigen receptors.^52^ ^53^ Unfortunately, the efficacy of NK cell-based therapies has been mixed in clinical trials, and the utility of NK cells against solid tumours is relatively unexplored. Here, we used a humanized mouse model equipped to support human NK cells to test the hypothesis that NK cells could be used to treat pancreatic adenocarcinoma (PDAC). We demonstrate that PDAC expresses ligands for both activating and inhibitory NK cells receptors, with both categories of ligands changing in response to immune pressure and inflammation. The changing nature of PDAC likely creates a challenge for immunotherapy but endorses the use of NK cells, whose expression of activating receptors differs throughout the repertoire, responding the continuously changing tumour landscape. We find that HLA I-driven inhibition dominates over activation of NK cells, interrupting the continuous targeting of these cancer cells, but that this can be overcome by selection of allogeneic, HLA-mismatched NK cells. The safety and efficacy of allogeneic NK therefore cells raises the possibility of donor selection to maximize reactivity by selecting NK cells refractory to the specific inhibitory ligands available on tumour cells.

PDAC tumours advance rapidly, with diagnosis most often at stage III or IV.^2^ While cure by surgery is possible, it is available to only a fraction of patients, and the risk for relapse remains high.^54^ In patients with PDAC, infiltration of NK cells into the solid tumour is rare, but the presence of circulating NK cells with increased production of IFN-γ is associated with improved patient outcomes.^55^ ^56^ It is only after the NK cells become dysregulated, decrease their production of inflammatory cytokines, and start producing the immunoregulatory cytokine IL-10 that they are no longer prognostically beneficial.^57–59^ As it is the immunosuppressive PDAC tumour microenvironment that drives this eventual dysregulation, we reasoned that insight into how NK cells may be inhibited by PDAC could help with therapeutic design to ensure that cellular immunotherapies have mechanisms to avoid this shut down of activation.

NK cells’ expression of activating and inhibitory receptors is heterogeneous, with co-expression of multiple receptor types common on the same subsets of NK cells that comprise an individual’s repertoire. We found, through genetic analyses and phenotypic confirmation, that PDAC tumours are dynamic in their expression of putative ligands for NK cells. With accumulating genetic mutations or inflammatory signaling, the ligands available for engagement with NK cell receptors are altered. We defined the natural cytotoxicity receptors (NCRs), NKp30 and NKp46, as well as DNAM-1 and NKG2D as putative contributors to the activation of NK cells against PDAC, but none were alone sufficient to predict NK anti-PDAC activity. These activating receptors are present throughout the NK cell repertoire, including on both educated and uneducated NK cells, and are likely to provide redundancy in the recognition of PDAC tumours. This echoes previous work that demonstrated that *ex vivo* expanded PDAC patient NK cells mediate killing of autologous PDAC tumours through NKG2D and DNAM-1, with some contribution of the NCRs.^60^ NKG2D has repeatedly be shown to be a key receptor in the anti-tumourgenicity of NK cells and driver of cytotoxicity against PDAC.^61^ Taken together, these findings endorse NKG2D and NCR as inclusion criteria in the design of NK cell-based immunotherapies.

When exposed to both immune pressure and IFN-γ induced inflammation, PDAC cells increased their expression of HLA I. During the double co-culture against AsPC-1 – where after 10 hours the primary PBMCs from the initial co-culture were removed and the tumours rinsed before a second round of PBMCs of either the same donor and a different donor were added for an additional five hours – NK cells were less responsive during the second round regardless of whether that specific donor’s NK cells have previously engaged with those same tumour cells. We inferred that this loss of activation was at least in part due to the upregulation of HLA I. This was not simply the result of tumour selective killing by the first round of NK cells because co-culturing PBMCs with tumour cells that had pre-treated with IFN-γ similarly led to decreased activation and concurrent upregulation of HLA I. By blocking HLA-ABC with antibodies, we confirmed that NK cells were more responsive when KIR-HLA inhibition was removed, revealing this interaction to be a major contributor to inhibition.

NK cells differentiate “self” from “non-self” by binding HLA I molecules in a process known as education.^12^ To the NK cell, HLA I molecules can be divided into groups based on their binding by specific KIR receptors at conserved public epitopes (“KIR ligands”). Unlike for T cells, where HLA allele-level matching is required, the interactions between KIR and KIR ligands are conserved between individuals.^28^ ^62^ Diversity within the HLA system at HLA-C and HLA-B, primarily, create binary opportunities for self/non-self discrimination.^62^ ^63^ Since almost all individuals’ KIR repertoires encode the major inhibitory KIR (*KIR2DL1, KIR2DL2/L3, KIR3DL1*), the combination of KIR and KIR ligands creates six combinations of NK cell “education” with only one represented by the “all ligands” moniker, where an inhibitory ligand is available for each of the three major KIR.^64^ Even among these “all ligands” patients, it may be possible to leverage KIR allelic variation or activating receptors in NK cell donors to effect an alloreactive impact in patients.^19^ ^65–67^ Hence, should selection of polyclonal populations from allogeneic sources be pursued, only a small number of “universal” donors would be needed to serve all patients’ potential combinations of KIR ligands, making off-the-shelf allogeneic therapy a realistic possibility.

It was largely the KIR3DL1^+^ subset of NK cells that co-expressed the activation receptors (NCRs, NKG2D) that responded to our tumour challenges when HLA inhibition was blocked, which prompted us to consider this population primarily in our *in vivo* studies. Education through KIR3DL1 may occur at least in part via *cis* interactions (i.e. on the NK cell itself). We have previously demonstrated that both donor and recipient HLA-Bw4 can contribute to NK cell education, and that even the “null” allele of KIR3DL1 (which is not presented on the cell surface) can be educated by “self” HLA-Bw4. ^19^ ^68^ Hence, even with adoptive allogeneic cell therapy, it is ostensibly possible to maintain NK cell education (and hence, continued function).

NK cells are known to have the advantage of allogeneic transplantability – they are not associated with graft-versus-host disease – opening the possibility for truly off-the-shelf and on-demand delivery of cellular therapies that can be adapted to a patient’s – and tumour’s – needs. NK cells are associated with lower toxicities, and may even be delivered in outpatient clinics.^24^ While these features endorse the use of NK cells as immunotherapy, work remains to define the ideal NK cell “platform” on which to build them. Careful consideration to the features of each tumour may provide clues toward NK cell subset selection, but given the heterogenous nature of PDAC (and other solid tumours), effective therapy will require agility and polyvalent capabilities for recognizing tumour cells to be effective. We demonstrate that it is the inhibitory interactions that dominate NK:PDAC outcomes, and that the availability of ligands for NK cell binding is highly dynamic. In this context, NK cells may be the ideal effectors, with already-diverse combinations of activating receptors and precise immunotherapy may require only removal of inhibitory interactions to enrich a population capable of continued activation. In light of the safety associated with allogeneic adoptive NK cell transfer, this may be achieved by selecting highly educated (activating receptor-equipped) populations of NK cells from KIR ligand-mismatched healthy donors.

## ACKNOWLEDGEMENTS

We are grateful to patient partners BP and KN, and to Dr. Lenny Schultz (The Jackson Labs) for his generous gift of HLA-Cw3 transgenic mice used in these experiments. SNL and RA are trainees in the Cancer Research Training Program of the Beatrice Hunter Cancer Research Institute. SNL is supported by a Canada Graduate Scholarship from the Canadian Institutes of Health Research and a Nova Scotia Graduate Scholarship through Dalhousie University. RA is supported by the Rick Salsman Pancreatic Cancer Studentship Fund through the Dalhousie Faculty of Medicine 2025 Graduate Studentship program. This work was supported by Craig’s Cause Pancreatic Cancer Society, and JD Irving and Innovation grants to JEB from the Canadian Cancer Society. Dalhousie University is located in Mi’kma’ki, the unceded and ancestral territory of the Mi’kmaq people.

