## Supplemental material for "Effective allogeneic natural killer cell therapy for pancreatic adenocarcinoma avails conserved activating receptors and evades HLA I-driven inhibition"

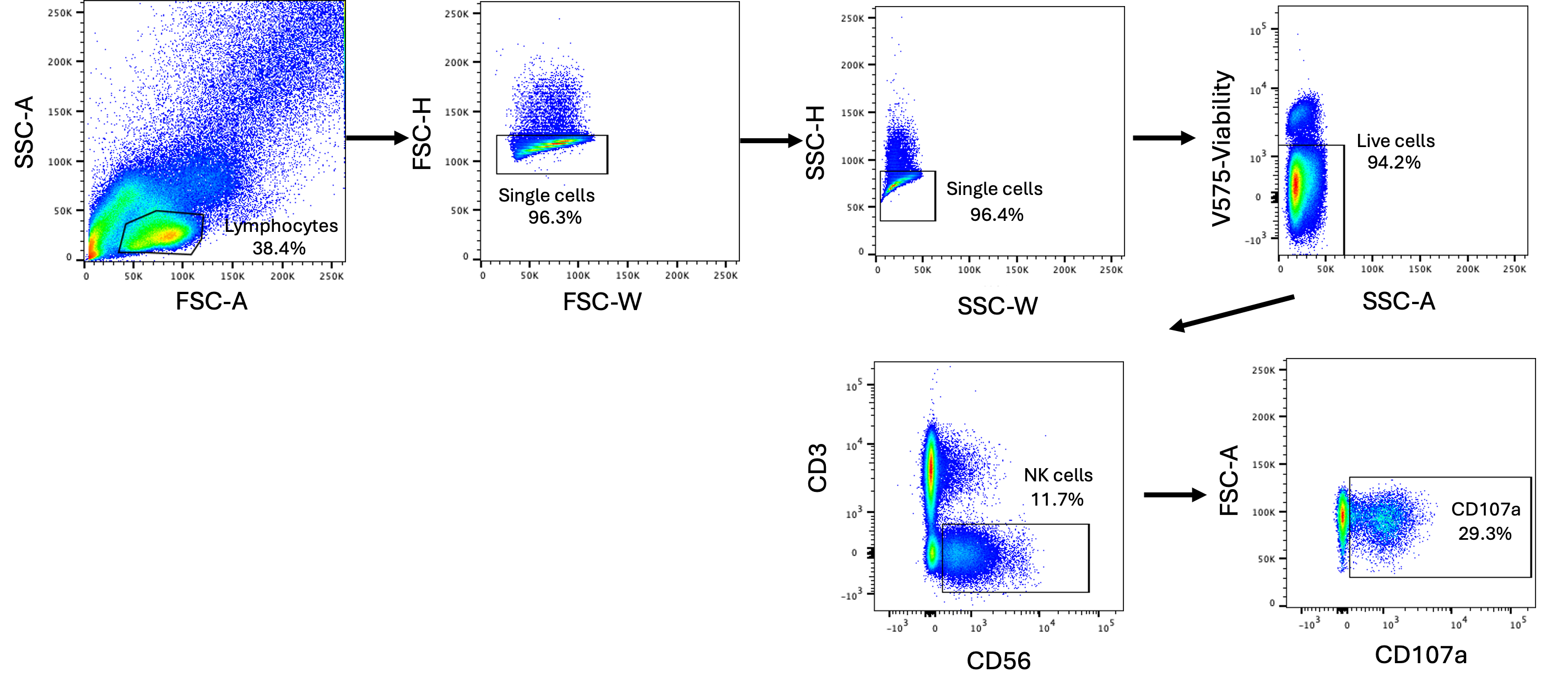

**Supplementary figure 1. Representative flow cytometry gating scheme for natural killer cells**

Lymphocytes were gated based on their size and complexity through the forward scatter area (FSC-A) and side scatter area (SSC-A), respectively. Duplicate cells were excluded by gating the single cells using the forward scatter height (FSC-H) and forward scatter width (FSC-W), followed by the side scatter height (SSC-H) and side scatter width (SSC-W). The cells were stained with the Viability Dye V575, which allowed for the exclusion of stained dead cells. NK cells were gated on as CD3^-^CD56^+^, followed by representative gating for CD107a.

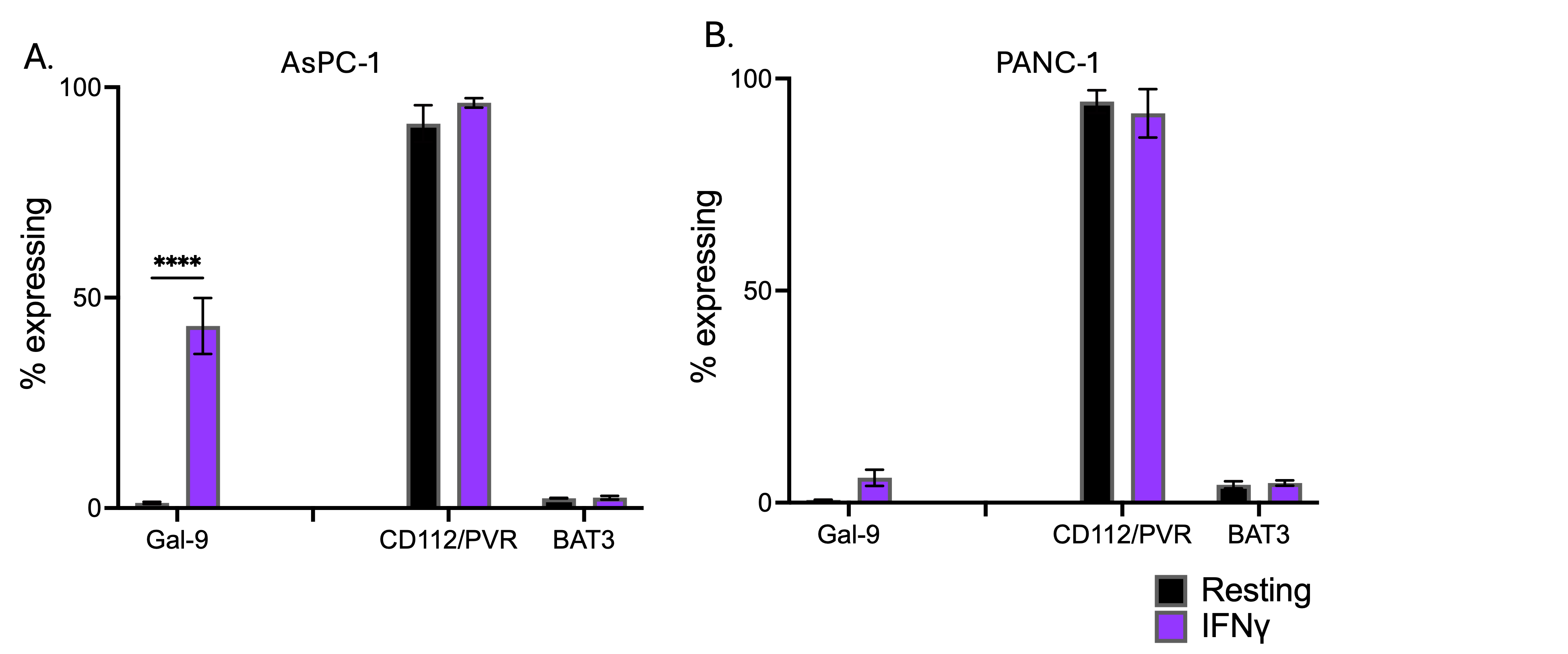

**Supplementary figure 2. Inflammatory signaling results in increased expression of inhibitory cancer ligands**

To simulate the more prolonged inflammatory environment associated with persistent NK cell activation, AsPC-1 and PANC-1 were treated with IFN-γ for three days. Tumour ligand expression was measured as percent of tumour cells expressing each marker. Ligand percent expression (A) AsPC-1 (B) PANC-1 (n=8). Statistics shown are mixed effects analysis with multiple comparisons ****=p<0.0001

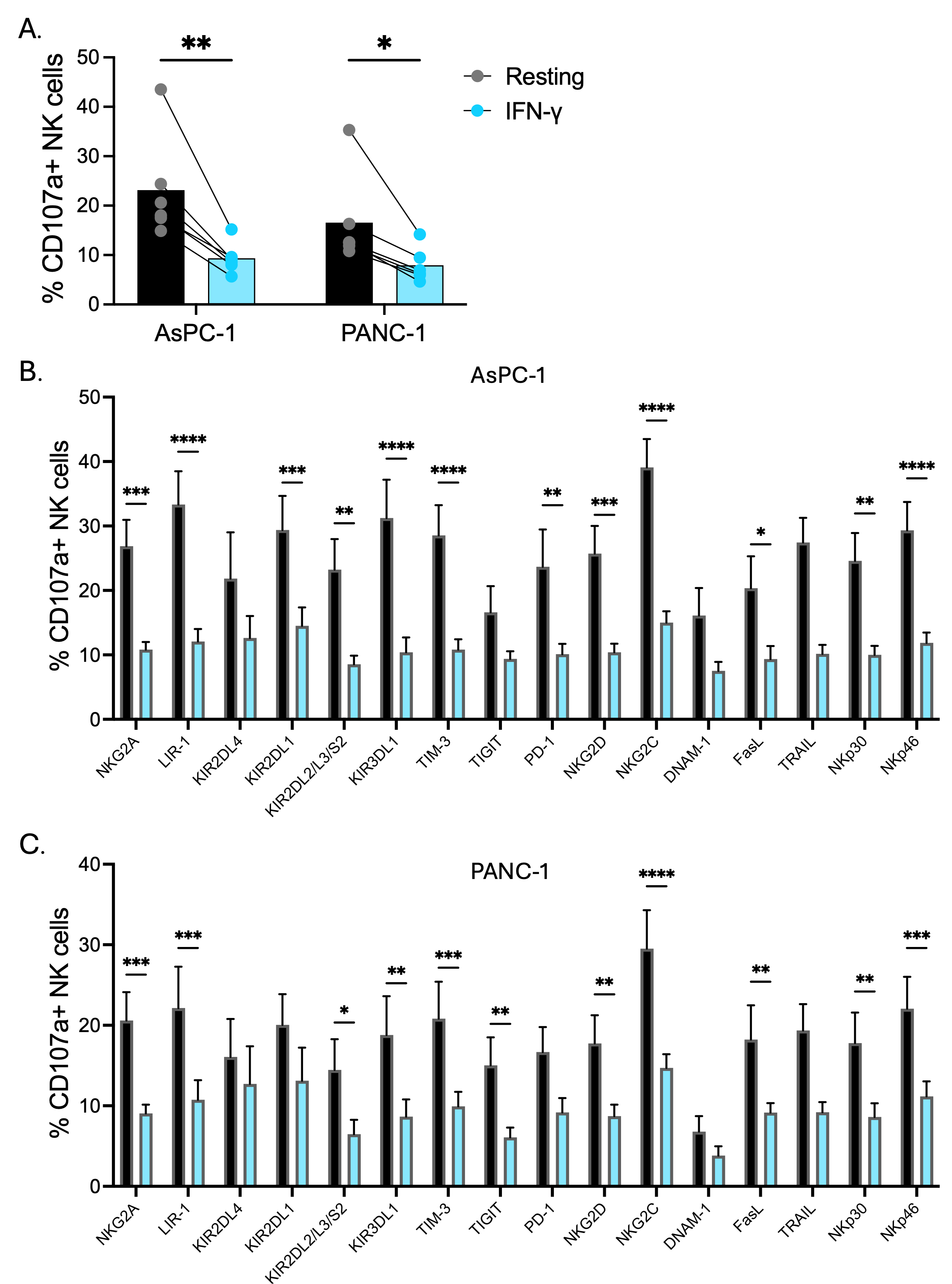

Supplementary figure 3. Treatment with IFN-γ results in decreased NK cell activation.

To simulate the inflammatory environment associated with NK cell activation, two PDAC cell lines, AsPC-1 and PANC-1, were treated with IFN-γ for three days prior to coculture with NK cells (3:1 E:T). CD107a (anti-LAMP1) was used to identify degranulating NK cells. To determine the relative representation of NK cell receptors among responding NK cells, NK cells were stained for flow cytometry after the co-culture. (A) The percentage of the total NK cell population responding to each cell line. Each dot represents a separate donor (n=6) (B) The percentage of responding NK cells expressing receptors for inhibition or activation after co-culture with AsPC-1 (C) The percentage of responding NK cells expressing receptors for inhibition or activation after co-culture with PANC-1. Statistics shown are two-ANOVA with multiple comparisons. *=p < 0.05; **=p<0.01; ***=p,0.001; ****=p<0.0001

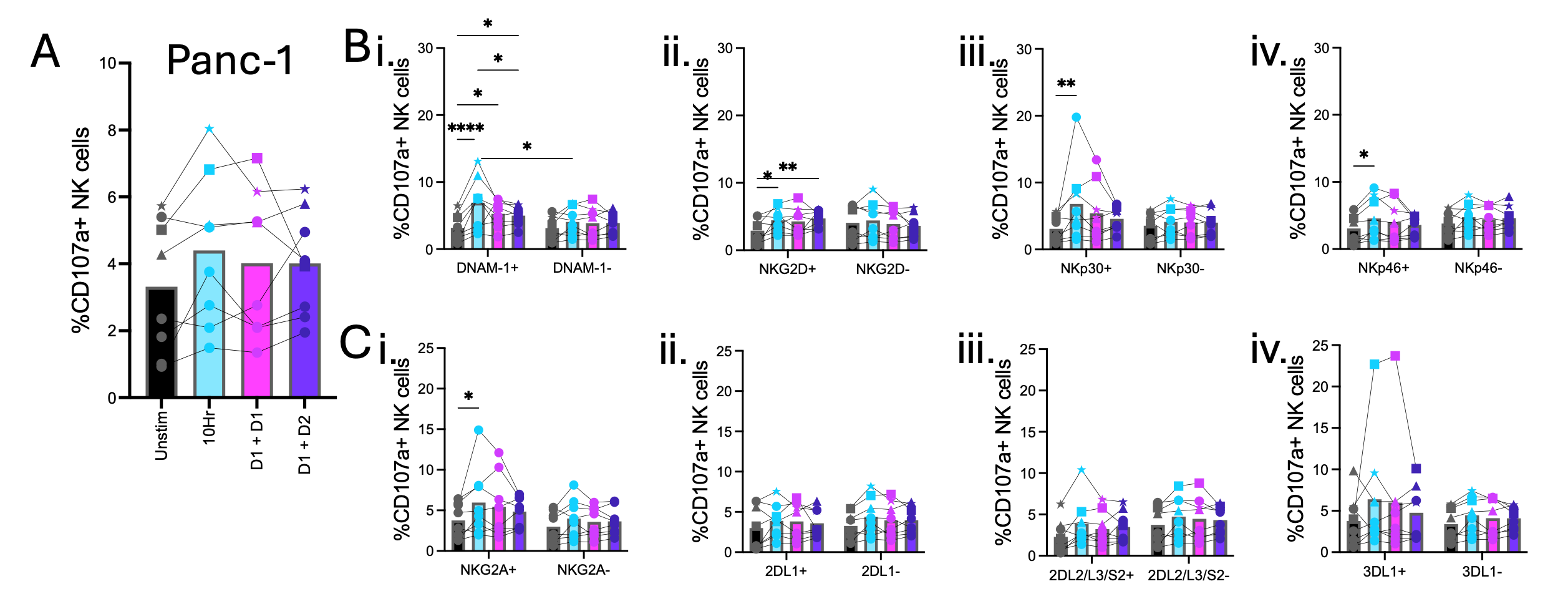

**Supplementary figure 4. Activation induced tumour ligand changes reduce responsiveness of primary NK cells against PANC-1.**

Exposure to activated immune cells can alter a tumour ligandome; to assess how these changes impacted the subsequent responsiveness of primary NK cells, PBMCs were co-culture with the PDAC cell line PANC-1 for 10-hour (10 Hr), then rinsed and re-co-cultured for 5-hour, with PBMCs of either the same donor (D1+D1) or a donor with a different educational background (D1 +D2) (A) Changes in total NK cell degranulation after 10hr or 10hr + 5hr co-culture with PANC-1 (B) Degranulation of activating receptor positive of negative populations (i. DNAM-1, ii. NKG2D, iii. NKp30, iv. NKp46). (n=8) Statistics shown are two-way ANOVA with multiple comparisons; *=p < 0.05; **=p<0.01; ***= p < 0.001; ****=p<0.0001

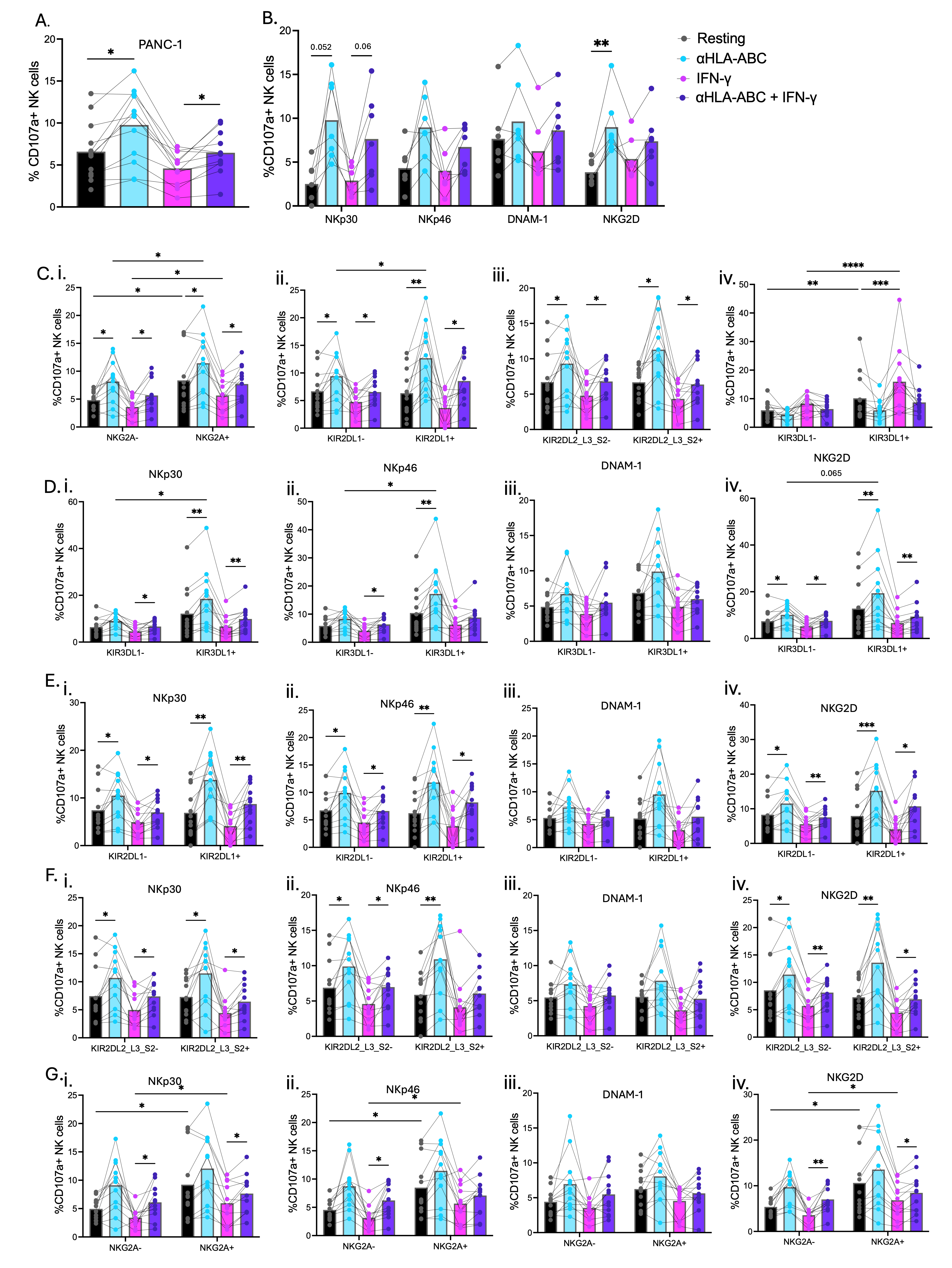

**Supplementary figure 5. HLA I inhibits NK cell reactivity against PDAC tumour cells.**

PANC-1 tumour cells were treated with IFN-γ or left untreated for three days, prior to co-culture with primary human NK cells (3:1 E:T) in the presence or absence of HLA-ABC blocking antibody (αHLA-ABC) to identify the impact of HLA-driven inhibition on NK reactivity against PDAC. (A) changes in total NK cell degranulation in response to PANC-1 treated with IFN-γ and/or HLA-ABC blocking after 5h of coculture. (B) Degranulation of single positive activating receptors (C) Degranulation of KIR- or KIR+ populations (i. NKG2A, ii. KIR2DL1, iii. KIR2DL2_L3_S2, or iv. KIR3DL1). (D-G) Activating receptor degranulation when co-expressing, or not (D) KIR3DL1 (i. NKp30, ii. NKp46, iii. DNAM-1, iv. NKG2D) (E) KIR2DL1 (i. NKp30, ii. NKp46, iii. DNAM-1, iv. NKG2D). (F) KIR2DL2/L3/S2 (i. NKp30, ii. NKp46, iii. DNAM-1, iv. NKG2D). (G) NKG2A (i. NKp30, ii. NKp46, iii. DNAM-1, iv. NKG2D). Lines join the responses of individual donors. (n=12); Statistics shown are two-way ANOVA with multiple comparisons; Statistics shown are two-way ANOVA with multiple comparisons; Statistics shown are two-way ANOVA with multiple comparisons; *=p<0.05; **=p<0.01; ***= p < 0.001; ****=p<0.0001

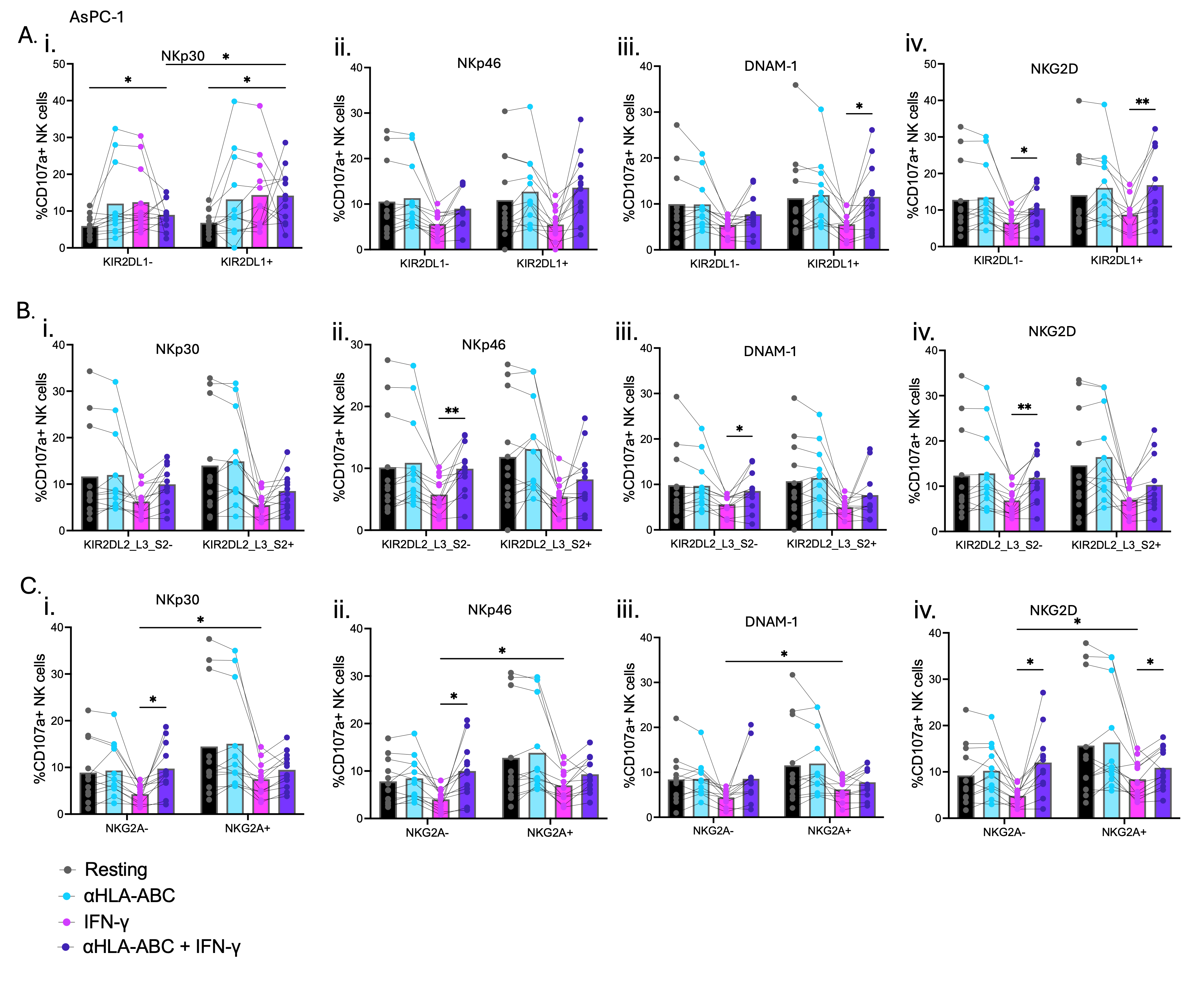

**Supplementary figure 6.** **Co**-**expression of KIR2DL1 and KIR2DL2/L3/S2 does not increase NK cell reactivity against AsPC-1.**

AsPC-1 tumour cells were treated with IFN-γ or left untreated for three days, prior to co-culture with primary human NK cells (3:1 E:T) in the presence or absence of HLA-ABC blocking antibody (αHLA-ABC) to identify the impact of HLA-driven inhibition on NK reactivity against PDAC. (A) Activating receptor degranulation when co-expressing, or not, KIR2DL1 (i. NKp30, ii. NKp46, iii. DNAM-1, iv. NKG2D). (B) Activating receptor degranulation when co-expressing, or not, KIR2DL2/L3/S2 (i. NKp30, ii. NKp46, iii. DNAM-1, iv. NKG2D). (C) Activating receptor degranulation when co-expressing, or not, NKG2A (i. NKp30, ii. NKp46, iii. DNAM-1, iv. NKG2D). Lines join the responses of individual donors. (n=12); Statistics shown are two-way ANOVA with multiple comparisons; *=p<0.05; **=p<0.01

Supplementary Table 1 Flow cytometry staining panel for human natural killer cells

| **Marker** | **Clone** | **Fluorochrome** | **Isotype** | **Company** | **Catalogue Number** |
| --- | --- | --- | --- | --- | --- |
| KIR3DL1 | DX9 | BV786 | Mouse | BD Biosciences | 742982 |
| CD39 | Tü66 | BV750 | Mouse | BD Biosciences | 747079 |
| CD73 | AD2 |  | Mouse | BD Biosciences | 747205 |
| TRAIL | RIK-2 | BV711 | Mouse | BD Biosciences | 743722 |
| CD3 | UCHT-1 | BV650 | Mouse | BD Biosciences | 563852 |
| NKp46 | 9-E2 | BV605 | Mouse | BD Biosciences | 743710 |
| Live/Dead |  | FVS570 |  | BD Biosciences | 564995 |
| TIGIT | 741182 | BV480 | Mouse | BD Biosciences | 747843 |
| NKG2C | 134591 | BV421 | Mouse | BD Biosciences | 748169 |
| PD-1 | EH12.1 | FITC | Mouse | R&D Systems | FAB10861G |
| KIR2DL1 | REA284 | PE-Cy7 | REA | Miltenyi Biotech | 130-120-447 |
| KIR2DL2/L3/S2 | GL183 | PE-Cy5.5 | Mouse | Beckman-Coulter | A66900 |
| TIM-3 | F38-2E2 | PE-Cy5 | Mouse | Biolegend | 345052 |
| NKG2D | 1D11 | PE-CF594 | Mouse | BD Biosciences | 562498 |
| KIR2DL4 | REA768 | PE | REA | Miltenyi Biotech | 130-112-465 |
| CD107a | H4A3 | APC-H7 | Mouse | BD Biosciences | 561343 |
| CD16 | 3G8 | AF700 | Mouse | BD Biosciences | 557920 |
| NKG2A | REA110 | APC | REA | Miltenyi Biotech | 130-113-563 |
| LIR-1 | GHI/75 | BUV805 | Mouse | BD Biosciences | 748454 |
| FasL | NOK-1 | BUV737 | Mouse | BD Biosciences | 748972 |
| NKp30 | p30-15 | BUV661 | Mouse | BD Biosciences | 750510 |
| NKp44 | p44-8 | BUV615 | Mouse | BD Biosciences | 752353 |
| CD8 | SK1 | BUV563 | Mouse | BD Biosciences | 741440 |
| DNAM-1 | DX11 | BUV496 | Mouse | BD Biosciences | 749935 |
| CD56 | NCAM16.2 | BUV395 | Mouse | BD Biosciences | 563554 |

Supplementary Table 2 Flow cytometry staining panel for tumour cell ligands in IFN-γ pre-treatment experiments

| **Marker** | **Clone** | **Fluorochrome** | **Isotype** | **Company** | **Catalogue number** |
| --- | --- | --- | --- | --- | --- |
| TRAIL-R2 | B-K29 | BV786 | Mouse | BD Biosciences | 745561 |
| TRAIL-R1 | S35-934 | BV711 | Mouse | BD Biosciences | 745424 |
| MIC A/B | 6D4 | BV650 | Mouse | BD Biosciences | 742325 |
| Live/Dead |  | FVS570 |  | BD Biosciences | 564995 |
| HLA-ABC | G46-2.6 | BV605 | Mouse | Biolegend | 311432 |
| **HLA-C** | **DT9** | **BV480** | **Mouse** | **BD Biosciences** | **747590** |
| PD-L1 | MIH1 | BV421 | Mouse | BD Biosciences | 563738 |
| PD-L2 | MIH18 |  | Mouse | BD Biosciences | 563842 |
| **B7-H6** | **1A5** | **BB790-P** | **Rat** | **BD Biosciences** | **Custom** |
| HLA-G | 87G | PerCP-Vio700 | Mouse | Miltenyi Biotech | 130-111-855 |
| HLA-E | 3D12 | PE-Cy7 | Mouse | Biolegend | 342608 |
| Fas | REA738 | PE-CF594 | REA | Miltenyi Biotech | 130-113-009 |
| **Gal-9** | **9M1-3** | **PE** | **Mouse** | **BD Biosciences** | **565890** |
| **CD112** | **11A8** | **APC** | **Mouse** | **BD Biosciences** | **567857** |
| HLA-Bw4 | REA274 | APC-H7 | REA | Miltenyi Biotech | 130-103-851 |
| ULBP-1 | 170818 | AF700 | Rabbit | R&D Systems | FAB1380N |
| CD276 | 7-517 | BUV661 | Mouse | BD Biosciences | 749896 |
| MHCII | TU39 | BUV563 | Mouse | BD Biosciences | 741379 |
| ULBP-2,5,6 | 165903 | BUV395 | Mouse | BD Biosciences | 748127 |

Supplementary Table 3 Flow cytometry staining panel for tumour cell ligands in double PBMC co-culture experiments

| **Marker** | **Clone** | **Fluorochrome** | **Isotype** | **Company** | **Catalogue number** |
| --- | --- | --- | --- | --- | --- |
| TRAIL-R2 | B-K29 | BV786 | Mouse | BD Biosciences | 745561 |
| TRAIL-R1 | S35-934 | BV711 | Mouse | BD Biosciences | 745424 |
| MIC A/B | 6D4 | BV650 | Mouse | BD Biosciences | 742325 |
| Live/Dead |  | FVS570 |  | BD Biosciences | 564995 |
| HLA-ABC | G46-2.6 | BV605 | Mouse | Biolegend | 311432 |
| PD-L1 | MIH1 | BV421 | Mouse | BD Biosciences | 563738 |
| PD-L2 | MIH18 |  | Mouse | BD Biosciences | 563842 |
| HLA-G | 87G | PerCP-Vio700 | Mouse | Miltenyi Biotech | 130-111-855 |
| HLA-E | 3D12 | PE-Cy7 | Mouse | Biolegend | 342608 |
| Fas | REA738 | PE-CF594 | REA | Miltenyi Biotech | 130-113-009 |
| **HLA-C** | **DT-9** | **PE** | **Mouse** | **BD Biosciences** | **566372** |
| **B7-H6** | **1A5** | **APC** | **Rat** | **BD Biosciences** | **566733** |
| HLA-Bw4 | REA274 | APC-H7 | REA | Miltenyi Biotech | 130-103-851 |
| ULBP-1 | 170818 | AF700 | Rabbit | R&D Systems | FAB1380N |
| CD276 | 7-517 | BUV661 | Mouse | BD Biosciences | 749896 |
| MHCII | TU39 | BUV563 | Mouse | BD Biosciences | 741379 |
| ULBP-2,5,6 | 165903 | BUV395 | Mouse | BD Biosciences | 748127 |

Supplementary Table 4: NK cell and immune ligand genes used in TCGA analysis

| **Gene Symbol** | **Protein Name (Abbreviated)** |
| --- | --- |
| *ADAM10* | ADAM10 |
| *ADAM17* | TACE |
| *B2M* | β2M |
| *BAG6* | BAG6 |
| *BMI1* | BMI-1 |
| *CADM1* | CADM1 |
| *CALR* | Calreticulin |
| *CCL19* | CCL19 |
| *CCL21* | CCL21 |
| *CD112* | Nectin-2 |
| *CD160* | CD160 |
| *CD1D* | CD1d |
| *CD2* | CD2 |
| *CD207* | Langerin |
| *CD226* | DNAM-1 |
| *CD244* | 2B4 |
| *CD273* | PD-L2 |
| *CD274* | PD-L1 |
| *CD276* | B7-H3 |
| *CD4* | CD4 |
| *CD48* | CD48 |
| *CD58* | LFA-3 |
| *CD59* | CD59 |
| *CD70* | CD70 |
| *CD72* | CD72 |
| *CD74* | CD74 |
| *CD84* | CD84 |
| *CD8A* | CD8α |
| *CD8B* | CD8β |
| *CD96* | CD96 |
| *CDH1* | E-cadherin |
| *CEACAM1* | CEACAM1 |
| *CFP* | C4b-binding protein |
| *CIITA* | Class II transactivator |
| *CREB1* | CREB1 |
| *CSF2* | GM-CSF |

**Supplementary Table 4. continued**

| **Gene Symbol** | **Protein Name (Abbreviated)** |
| --- | --- |
| *CTSL* | Cathepsin L |
| *CTSS* | Cathepsin S |
| *CX3CL1* | CX3CL1 |
| *CXCL10* | CXCL10 |
| *CXCL11* | CXCL11 |
| *CXCL12* | CXCL12 |
| *CXCL2* | CXCL2 |
| *CXCL8* | IL-8 |
| *CXCL9* | CXCL9 |
| *ERAP1* | ERAP1 |
| *FAS* | Fas (CD95) |
| *FCER1G* | FcεRIγ |
| *FCGR1A* | FcγRI |
| *FCGR3A* | FcγRIIIa |
| *FOXO1* | FOXO1 |
| *GATA2* | GATA2 |
| *GZMB* | Granzyme B |
| *HFE* | HFE |
| *HLA-A* | HLA-A |
| *HLA-B* | HLA-B |
| *HLA-C* | HLA-C |
| *HLA-DMA* | HLA-DMA |
| *HLA-DMB* | HLA-DMB |
| *HLA-DOA* | HLA-DOA |
| *HLA-DOB* | HLA-DOB |
| *HLA-DPA1* | HLA-DPA1 |
| *HLA-DPB1* | HLA-DPB1 |
| *HLA-DQA1* | HLA-DQA1 |
| *HLA-DQA2* | HLA-DQA2 |
| *HLA-DQB1* | HLA-DQB1 |
| *HLA-DRA* | HLA-DRA |
| *HLA-DRB1* | HLA-DRB1 |
| *HLA-DRB3* | HLA-DRB3 |
| *HLA-DRB4* | HLA-DRB4 |
| *HLA-DRB5* | HLA-DRB5 |
| *HLA-E* | HLA-E |

**Supplementary Table 4. continued**

| **Gene Symbol** | **Protein Name (Abbreviated)** |
| --- | --- |
| *HLA-F* | HLA-F |
| *HLA-G* | HLA-G |
| *HMGB1* | High mobility group box 1 |
| *HSP90AA1* | Heat shock protein 90 alpha 1 |
| *HSP90AB1* | Heat shock protein 90 alpha B1 |
| *HSPA1A* | Heat shock protein 70kDa 1A |
| *HSPA1B* | Heat shock protein 70kDa 1B |
| *HSPA1L* | Heat shock protein 70kDa 1L |
| *HSPA2* | Heat shock protein 70kDa 2 |
| *HSPA4* | Heat shock protein 70kDa 4 |
| *HSPA5* | Heat shock protein 70kDa 5 |
| *HSPA6* | Heat shock protein 70kDa 6 |
| *HSPA8* | Heat shock protein 70kDa 8 |
| *HSPG2* | Heparan sulfate proteoglycan 2 (perlecan) |
| *IFI30* | IFI30 |
| *IFNG* | IFN-γ |
| *IL10* | IL-10 |
| *IL12A* | IL-12p35 |
| *IL12B* | IL-12p40 |
| *IL15* | IL-15 |
| *IL18* | IL-18 |
| *IL2* | IL-2 |
| *IL21* | IL-21 |
| *IL23A* | IL-23 |
| *IL37* | IL-37 |
| *ITGB5* | Integrin β5 |
| *KIR2DL1* | KIR2DL1 |
| *KIR2DL2* | KIR2DL2 |
| *KIR2DL3* | KIR2DL3 |
| *KIR2DL4* | KIR2DL4 |
| *KIR2DL5A* | KIR2DL5A |
| *KIR2DS1* | KIR2DS1 |
| *KIR2DS2* | KIR2DS2 |
| *KIR2DS3* | KIR2DS3 |
| *KIR2DS4* | KIR2DS4 |
| *KIR2DS5* | KIR2DS5 |

**Supplementary Table 4. continued**

| **Gene Symbol** | **Protein Name (Abbreviated)** |
| --- | --- |
| *KIR3DL1* | KIR3DL1 |
| *KIR3DL2* | KIR3DL2 |
| *KIR3DL3* | KIR3DL3 |
| *KIR3DS1* | KIR3DS1 |
| *KLRC1* | NKG2A |
| *KLRC2* | NKG2C |
| *KLRC3* | NKG2E |
| *KLRC4* | NKG2F |
| *KLRD1* | CD94 |
| *KMT2E* | MLL5 (Mixed-Lineage Leukemia 5) |
| *LGALS9* | Galectin-9 |
| *LGMN* | Legumain |
| *LILRB1* | ILT3 |
| *LY9* | CD229 (Lymphocyte Antigen 9) |
| *MICA* | MICA |
| *MICB* | MICB |
| *MYC* | c-Myc |
| *NCF1* | p47phox |
| *NCR3LG1* | B7-H6 |
| *NECTIN2* | CD112 |
| *NECTIN3* | Nectin-3 |
| *NECTIN4* | Nectin-4 |
| *NFYA* | NF-YA |
| *NFYB* | NF-YB |
| *NFYC* | NF-YC |
| *NID1* | Nidogen-1 |
| *KLRK1* | NKG2D |
| *NCR3* | NCR3 |
| *NCR2* | NCR2 |
| *NCR1* | NCR1 |
| *KLRF2* | NKp46 |
| *KLRF1* | NKp44 |
| *PCNA* | Proliferating cell nuclear antigen |
| *PDCD1LG2* | PD-L2 |
| *PDIA3* | PDIA3 |
| *PFR1* | PFR1 |

**Supplementary Table 4. continued**

| **Gene Symbol** | **Protein Name (Abbreviated)** |
| --- | --- |
| *PSMA2* | Proteasome subunit alpha type-2 |
| *PSMB1* | Proteasome subunit beta type-1 |
| *PSMB10* | Proteasome subunit beta type-10 |
| *PSMB2* | Proteasome subunit beta type-2 |
| *PSMB6* | Proteasome subunit beta type-6 |
| *PSMB7* | Proteasome subunit beta type-7 |
| *PSMB8* | Proteasome subunit beta type-8 |
| *PSMB9* | Proteasome subunit beta type-9 |
| *PSMC4* | Proteasome 26S subunit ATPase 4 |
| *PSMC5* | Proteasome 26S subunit ATPase 5 |
| *PSMD1* | Proteasome 26S subunit, non-ATPase 1 |
| *PSMD13* | Proteasome 26S subunit, non-ATPase 13 |
| *PSMD2* | Proteasome 26S subunit, non-ATPase 2 |
| *PSMD4* | Proteasome 26S subunit, non-ATPase 4 |
| *PSMD5* | Proteasome 26S subunit, non-ATPase 5 |
| *PSMD7* | Proteasome 26S subunit, non-ATPase 7 |
| *PSME1* | Proteasome activator subunit 1 |
| *PSME2* | Proteasome activator subunit 2 |
| *PSME3* | Proteasome activator subunit 3 |
| *PVR* | Poliovirus receptor |
| *RAET1E* | RAET1E |
| *RAET1G* | RAET1G |
| *RAET1L* | RAET1L |
| *RFX5* | Regulatory factor X5 |
| *RFXANK* | RFX ankyrin repeat protein |
| *RFXAP* | RFX-associated protein |
| *SLAMF1* | SLAMF1 |
| *SLAMF6* | SLAMF6 |
| *SLAMF7* | SLAMF7 |
| *TAP1* | TAP1 (Transporter associated with antigen processing 1) |
| *TAP2* | TAP2 (Transporter associated with antigen processing 2) |
| *TAPBP* | TAP-binding protein (TAPBP) |
| *TGFB1* | TGF-β1 |
| *TGFB2* | TGF-β2 |
| *TIGIT* | TIGIT |

**Supplementary Table 4. continued**

| **Gene Symbol** | **Protein Name (Abbreviated)** |
| --- | --- |
| *TNF* | Tumor necrosis factor |
| *TNFRSF10A* | TRAIL receptor 1 (DR4) |
| *TNFRSF10B* | TRAIL receptor 2 (DR5) |
| *ULBP1* | ULBP1 |
| *ULBP2* | ULBP2 |
| *ULBP3* | ULBP3 |
